# Focal cortical dysplasia type II-dependent maladaptive myelination in the human frontal lobe

**DOI:** 10.1101/2024.03.02.582894

**Authors:** Catharina Donkels, Susanne Huber, Theo Demerath, Christian Scheiwe, Mukesch J. Shah, Marcel Heers, Horst Urbach, Andreas Schulze-Bonhage, Marco Prinz, Ute Häussler, Andreas Vlachos, Jürgen Beck, Julia M. Nakagawa, Carola A. Haas

## Abstract

Focal cortical dysplasias (FCDs) are local malformations of the human neocortex and a leading cause of intractable epilepsy. FCDs are classified into different subtypes including FCD IIa and IIb, characterized by a blurred gray-white matter boundary or a transmantle sign indicating abnormal white matter myelination. Recently, we have shown that myelination is also compromised in the gray matter of FCD IIa of the temporal lobe. Since myelination is key for brain function which is imbalanced in epilepsy, in the current study we investigated myelination in the gray matter of FCD IIa and IIb from the frontal lobe. We found that in particular FCD IIb showed myelination disturbances such as increased numbers of myelinating oligodendrocytes (OLs) and an irregular and disorganized myelination pattern covering an enlarged area in comparison to FCD IIa and controls. Interestingly, both FCD types presented with larger axon diameters when compared to controls. A significant correlation of axon diameter and myelin sheath thickness was found for FCD IIb and controls, whereas in FCD IIa large caliber axons were less myelinated. On the level of gene expression, FCD IIb presented with a significant up-regulation of myelin-associated mRNA synthesis in comparison to FCD IIa and by enhanced binding-capacities of the transcription factor MYRF to promoters of myelin-associated genes reflecting the need for more myelin due to increased axon diameters. These data show that FCD IIa and IIb are characterized by divergent signs of maladaptive myelination which may contribute to the epileptic phenotype.

**Main points:**

- In the gray matter of the frontal lobe, FCD IIa and FCD IIb are characterized by divergent signs of maladaptive myelination.
- FCD IIa presents with an ordinary radial fiber pattern, but with a reduced thickness of the myelin sheath around large diameter axons and with an attenuation of the myelin synthesis machinery.
- FCD IIb is characterized by an irregular and disorganized myelin fiber pattern, a higher density of myelinating oligodendrocytes and an elevated transcriptional turnover of myelin-associated genes.

## Introduction

Focal cortical dysplasias (FCDs) are a frequent cause of pharmcoresistant focal epilepsy, especially in children (Palmini et al., 1991, Fauser et al., 2004, Fauser et al., 2006). Surgical resection of the lesion remains often the only therapeutic option, resulting in a seizure-free clinical outcome in many cases (Zhao et al., 2020, Willard et al., 2022). FCDs comprise a heterogeneous group of cortical malformations morphologically characterized by disturbed lamination and cytoarchitecture including radial and/or horizontal alterations in lamination (FCD Ia or Ib) and the occurrence of dysmorphic neurons (FCD IIa) and/or balloon cells (FCD IIb) (Blümcke et al., 2011, Najm et al., 2022). Genetically, sporadic somatic mutations in the *MTOR* (mammalian target of rapamycin) gene and in mTOR pathway-related genes have been identified in FCD IIa and IIb cases leading to hyperactivation of mTOR signaling (Abdijadid et al., 2015, Lim et al., 2015, Baldassari et al., 2019, Nguyen et al., 2019, Chung et al., 2023). Recently, mutations in the GATOR1 complex, a GTPase-activating protein (GAP) complex that regulates TORC1 signaling by interacting with the Rag GTPase, have been found exclusively in FCD IIa of the frontal lobe (Honke et al., 2023), pointing out pathophysiological differences between FCD IIa and IIb. Accordingly, in the MRI many FCD IIa cases present with a blurring of the gray–white matter boundary (Blümcke et al., 2011, Mühlebner et al., 2012, Najm et al., 2022) , whereas FCD IIb shows a funnel-shaped formation through the white matter (WM) (transmantle sign) (Urbach et al., 2002, Kimura et al., 2019), which is considered as abnormal WM myelination.

Correct myelination is very important for proper brain function and a highly dynamic process involved in memory consolidation in the adult brain (Steadman et al., 2020). There is recent evidence, that maladaptive myelination plays also a role in epileptic conditions (Knowles et al., 2022, De Curtis et al., 2021). Indeed, recent reports show a loss of oligodendrocytes (OL) and myelination capacity in the white matter of FCD lesions (Mühlebner et al., 2012, Shepherd et al., 2013, Scholl et al., 2017, Gruber et al., 2021). Likewise, deficits in myelination were found in the gray matter of temporal lobe FCDs (Donkels et al., 2017), where impaired proliferation of oligodendrocyte precursor cells (OPC) is part of the FCD pathology (Donkels et al., 2020).

During the differentiation of OPCs into myelinating OLs, a number of transcription factors (TFs) are sequentially expressed to specify the OL fate (Emery and Lu, 2015, Sock and Wegner, 2021). Depending on the differentiation stage, these TFs inhibit or stimulate the expression of myelin-associated genes (Emery, 2010). Oligodendrocyte transcription factor 2 (OLIG2) is a prominent TF that initiates the differentiation of OPCs into myelinating OLs and is expressed across the entire lineage but with decreasing amounts toward the termination of differentiation (Mei et al., 2013, Meijer et al., 2014). A key factor for the completion of OL differentiation is the myelin regulatory factor (MYRF) (Emery et al., 2009, Bujalka et al., 2013), induced during terminal OL differentiation. MYRF positively regulates the transcription of myelin-associated genes encoding myelin basic protein (MBP), 2′, 3′-cyclic nucleotide-3′-phosphodiesterase (CNPase), myelin associated glycoprotein (MAG) and myelin oligodendrocyte glycoprotein (MOG) by binding to their promoters (Emery et al., 2009, Bujalka et al., 2013).

In the current study, we investigated potential alterations of myelination in the gray matter of FCD IIa and IIb cases localized in the human frontal lobe. We studied the distribution of OLs, the myelination pattern, the ultrastructure of axons and myelin sheaths and the transcriptional regulation of myelin-associated genes. We present evidence for an FCD-type specific dysregulation of myelination: FCD IIa is characterized by thinner myelin sheaths of large diameter axons and an attenuation of the myelin synthesis machinery. In contrast, FCD IIb presents with an irregular myelination pattern, with thicker axon diameters and myelin sheaths and an elevated transcriptional turnover of myelin-associated genes in comparison to FCD IIa.

## Methods

### Patient Selection

A total of 25 surgically resected frontal lobe specimens from patients with FCD IIa (n=9) [(age at surgery (years): mean: 16.78, range: 6 – 38), (epilepsy duration (years): mean: 10.24, range: 3 – 34)], FCD IIb (n=9) [(age at surgery (years): mean: 33.67, range: 12 – 48), (epilepsy duration (years): mean: 24.33, range: 9 – 41)] and epileptic but non-dysplastic controls (n=7) [approach tissue from tumor resections (WHO grade I) or non-FCD lesions] [(age at surgery (years): mean: 18.54, range: 6.1 – 30), (epilepsy duration (years): mean: 10.97, range: 4.1 – 23)] were included in this study. The inclusion of epileptic controls in our study was intentional to exclude epilepsy-related features and to focus on FCD-specific pathological alterations. Individual patients were color-coded to facilitate their identification in the different assays (able 1).

All cases had undergone neurosurgical interventions due to pharmacoresistant epilepsy or low-grade tumors and removal of cortical tissue was clinically warranted to achieve seizure or tumor control. Pre-surgical assessment included the documentation of a detailed history, neurological examination, neuropsychological testing, MRI scanning and non-invasive/invasive, long-term video EEG monitoring. Informed consent was obtained from all individuals included in the study. All procedures received prior approval by the institutional review board (Ethics Committee, Medical Center -University of Freiburg) and were in accordance with the 1964 Helsinki declaration and its later amendments.

For histopathological diagnosis, all specimens were classified on paraffin sections by the Institute of Neuropathology, Medical Center – University of Freiburg) according to Najm et al., 2022.

### Tissue Preparation

Brain tissue was collected immediately after resection in ice-cold 0.1M phosphate buffer (PB), pH 7.4. For immunohistochemistry (IHC) and *in situ* hybridization histochemistry (ISHH), slices (∼5 mm) were cut perpendicular to the brain’s surface, fixed in 4% paraformaldehyde (PFA) [w/v] in PB (24 - 48h, 4°C), cryoprotected in 30% sucrose in PB (24 - 48h, 4°C) and finally stored at -80°C until further processing. For IHC and ISHH, cryostat sections (50 μm, orthogonal to the cortical surface) were prepared, collected in tissue culture dishes, and either rinsed in PB (IHC) or in 2 X SSC (1 X SSC = 0.15 M NaCl, 0.015 M sodium citrate, pH 7.0) for ISHH.

For electron microscopy (EM), PFA-fixed frozen tissue was cut perpendicularly to the pia in a cryostat (60 µm) (Leica CM3050 S) and slices were post-fixed in 4% PFA/2.0% glutaraldehyde (GA) in PB (approx. 24h, 4°C) as described previously (Donkels et al., 2017).

For molecular biological techniques the tissue was immediately frozen at -80°C and stored until further processing.

### Immunohistochemistry

For NeuN, SMI32 and Vimentin immunolabeling a free-floating protocol was applied. Tissue slices were pretreated in 10% normal serum (NS) and 0.25% Triton X-100 for 30 min at RT and subsequently incubated in 0.1% Triton X-100 and 1% NS in PB with the following primary antibodies: mouse monoclonal anti-NeuN (1:1000; Millipore, #MAB377), anti-SMI32 (specific for non-phosphorylated neurofilament H, 1:1000; Biolegend #801701) and anti-Vimentin (1:1000; Dako #M0725) at room temperature (RT) overnight. The antibody binding was visualized by incubation with secondary antibodies conjugated with Cyanine (Cy)-3 (1:400; Jackson ImmunoResearch Laboratories) for 2h. Sections were mounted and coverslipped with Immu-Mount (Thermo Scientific).

A modified IHC protocol published by Moreno-Jiménez et al., 2019 was used for the CNPase staining. Tissue sections were incubated in a 0.5% sodium borohydride (NaBH_4_, Sigma-Aldrich) solution for 30 min at RT. Next, a heat-mediated 10 mM citrate buffer (pH 6.0) antigen retrieval step was performed by boiling the tissue sections in a water bath for 20 min followed by rinsing in 0.1 M PB and pretreatment in 0.5% bovine serum albumin (BSA) and 0.5% Triton for 30 min at RT. Incubation with the CNPase antibody (1:100, Cell Signaling Technology #5664S) was performed under gentle shaking at 4 °C for 5 days followed by treatment with the Cy-3-labelled secondary antibody (1:400; Jackson ImmunoResearch Laboratories) for 24h at 4°C under constant shaking. Additionally, all sections were counterstained for 10 min with 4,6-diamidino-2-phenylindole (DAPI,1:10,000) to label nuclei. Afterwards, slices were washed three times with PB, mounted on gelatinized microscope slides and treated with an Autofluorescence Eliminator reagent (EMD Millipore) for 5 min following the manufacturer’s instructions. Finally, the slices were washed three times with 70% ethanol and coverslipped in aqueous mounting medium (ImmuMount; Thermo Scientific).

### In situ Hybridization Histochemistry (ISHH)

*Mbp* mRNA expression was investigated by ISHH using digoxigenin (DIG)-labeled anti-sense riboprobes, generated from a pBluescript KS II^+^ vector containing a 1.2 kb *mbp* cDNA as previously described (Donkels et al., 2017).

In brief, cryostat sections (50 µm) were hybridized with a DIG-labeled anti-sense *mbp* cRNA riboprobe (100 ng/ml) at 55°C overnight followed by several stringent washing steps at 65°C. DIG-labeled hybrids were detected with an anti-DIG antibody from sheep conjugated with alkaline phosphatase as recommended by the manufacturer (Roche) by incubating the slice overnight at 4°C. For signal detection, nitro blue tetrazolium chloride and 5-bromo-4-chloro-3-indolylphosphate were applied and the reaction was developed in the dark. Finally, sections were coverslipped and mounted in Kaiser’s gelatin.

### Light microscopy and quantitative analyses

Fluorescence and bright field microscopy were performed with an *AxioImager2* microscope (Zeiss) using a 10x (NA 0.45) or 20x (NA 0.75) objective, (both Zeiss). Photomicrographs were taken as tile images with a digital camera (MR605) for fluorescence, with MR503 for bright field (both Zeiss), and processed with *Zen* software (Zeiss).

Confocal images were taken using the *Fluoview* FV10i (Olympus) and a 63x oil objective applying the following parameters: image size 1024×1024 pixel, sensitivity 50% and laser power 20% for Cy3 and DAPI. The numerical aperture was selected 1, stacks were taken with 0.5 µm step sizes.

For the quantification *of mbp* mRNA-positive cells, bright field images of three sections per patient were imported into *ImageJ* software. In each image, two regions of interest (ROIs) covering all six cortical layers were defined and within each ROI all digoxigenin-labeled cells were manually counted using the integrated “cell counter” plugin (National Institutes of Health).

For measurement of the myelinated area, images of CNPase/DAPI-labeled slices (two/patient) of the three were imported into the Fiji ImageJ software. In each section, two regions of interest (ROIs) were defined: ROI 1 (entire cortical gray matter) and ROI 2 (area of CNPase-positive fiber strands). The proportion of the myelinated cortex was calculated (myelinated area mm^2^/whole area mm^2^ X 100%) (Van Tilborg et al., 2017).

For quantitative determination of OL and pre-OL densities, confocal image stacks of CNPase-immunolabled sections were imported into *Image J*. OL and pre-OL cell bodies were counted in three slices per patient. In each slice, three ROIs in layers II/III and V/VI (18 ROIs/patient) were manually counted by starting from the first image, where cells appeared and stopped at the last countable image. Cell numbers were related to the volume of the counted stacks and given as a mean over all ROIs per patient (cell number/mm^3^).

### Electron Microscopy

For electron microscopy, tissue slices were cut perpendicularly to the pia from the fixed, frozen block (60µm) and post-fixed in the presence of 4% PFA and 2% glutaraldehyde for 24h at 4°C as described before (Donkels et al., 2020). For each patient, representative sections were stained with 1% cresyl violet to visualize the cortical layers, required for orientation before embedding. The other slices were osmicated for 2 h with 1% osmium tetroxide in 6.86% sucrose in PB, subsequently contrasted with 1% uranyl acetate in 70% ethanol overnight at 4°C and dehydrated in graded ethanol. The cortical gray matter was flat-embedded in epoxy resin (Durcupan ACM, Sigma-Aldrich), which was polymerized overnight. Ultrathin sections (60 nm) in the horizontal plane were prepared with an ultramicrotome (LEICA UC6) and collected on copper grids. Sections were analyzed with a TEM LEO 906E (Zeiss). Electron photomicrographs of layers V/VI were taken with the CCD-Camera “sharp eye” (Tröndle) and visualized with ISP software (Tröndle).

For quantitative analysis images were imported into *ImageSP* (TRS & SYSPROG). With this software, the outer and the inner diameter of myelinated axons were determined and the myelin sheath thickness and the g-ratio (the ratio of axon to total diameter) were calculated (Rushton, 1951).

### RNA Isolation, Reverse Transcription and Real-Time RT-qPCR Analysis

The isolation of total RNA, reverse transcription, and real-time qPCR was performed as described earlier (Donkels et al., 2017). In brief, frozen specimens were thawed on ice and the white matter was carefully removed with a scalpel in *RNAlater* (Qiagen). The total RNA was isolated from 30 mg cortical gray matter using the *RNeasy Mini kit* (Qiagen) according to the manufacturer’s guidelines. Afterwards, a reverse transcription (RT) step was carried out with 1µg total RNA per reaction using the *First Strand cDNA Synthesis Kit* (ThermoFisher Scientific).

The expression of mRNAs was quantified by real-time RT-qPCR on a *CFX Connect^TM^* Real-Time PCR System (Bio-Rad Laboratories) in the presence of *SsoAdvanced universal SYBR Green Mix* (Bio-Rad Laboratories). The following human-specific primer pairs were used at 70 nM: *s12* (mitochondrial ribosomal protein S12) (forward (f): 5’-AGTTGGTGGAGGCCCTTTGT-3’, reverse (r): 5’-AGGCCTACCCATTCTCCTAGTTTC-3’), *mbp* (f: 5’-AATTACTCACCGAGACACAC-3’, r: 5’-GAAGGATATGGTCAGAAGAG-3’), *mag* (f: 5’-CAACTGAGTCCAAGTTGTCT-3’, r: 5’-ATTCTGTACAGCGAGTGAGT-3’), *mog* (f: 5’-CTTCGAGCAGAGATAGAGAA-3’, r: 5’-TCGATGTAGCCAGTTGTAG-3’), *myrf* (f: 5’-CTACCACATCCCTGTCAGT-3’, r: 5’-GGAGGAGGAGTTCATCTG-3’), *cnp* (f: 5’-ACACACAGCTTCTAGTGATTT-3’, r: 5’-ACAGAGAAGCGAGTTCAG-3’), *olig2* (f: 5’-AATCTCAATATCTGGGTC AA-3’, r: 5’-GGAACATCCACAGATTTATT-3’), *sip1* (Smad interacting protein 1) (f: 5’-AGCCTCTGTAGATGGTCCAGAAGAA-3’, r: 5’-GCGGTCTGGATCGTGGCTTC-3’).

The PCR conditions were chosen as published before (Donkels et al., 2017). The resulting Ct (cycle threshold) values (obtained by the CFX Manager^TM^ Software, Bio-Rad Laboratories) were used to calculate relative expression levels for genes of interest by normalization to the housekeeping gene S12.

### Chromatin immunoprecipitation (ChIP)

The ChIP assays were performed using the *iDeal ChIP-seq* kit for transcription factors (Diagenode) with fresh-frozen FCD and control tissue (30 mg per IP) according to manufacturer’s guidelines. Firstly, the gray matter was manually dissected and homogenized with 1% formaldehyde to cross-link genomic DNA and proteins for 15 min at RT. The chromatin lysate was sonicated and chromatin was fragmented to a size of approx. 500-1000 base pairs using the *Bioruptor® Pico* (Diagenode) by running 10 cycles with settings 30 sec. on, 30 sec. off per cycle. To reduce non-specific binding, we included a preclearing step by incubating the chromatin lysate with *DiaMag* Protein A-coated magnetic beads (Diagenode) at 4°C for 1.5 - 2 hours. Next, polyclonal rabbit MYRF (5 µg, Millipore), polyclonal goat OLIG2 (5 µg, R&D) antibodies and non-specific polyclonal rabbit IgG (1 µg, Diagenode, as negative control) were pre-incubated for 4h at 4°C with Protein A-coated magnetic beads followed by addition of the precleared chromatin overnight at 4°C under constant rotation. Afterwards, the magnetic beads were stringently washed, the precipitated chromatin was eluted from the beads and the crosslinking of the DNA/transcription factor complex was reversed by incubating the eluate at 65°C overnight. Subsequently, the DNA was purified using *IPure beads v2* (Diagenode) and stored at -20°C until further use.

ChIP specificity was verified by real-time qPCR for known transcription factor binding sites (e.g., *MAG*; *MOG*, *MBP*). The PCR primers were designed around predicted transcription factor binding sites (BS), which were identified by using the *MatInspector* tool form *Genomatix* software. Using this tool, we detected the following transcription factor BS sequences: *MBP* (MYRF BS): 5’-TTTCCTGGTACAT-3’, *SIP 1* (OLIG2 BS): 5’-AGGCCAGCTGTTTCA-3’, *CNP* (OLIG2 BS): 5’-AGCCCCAGCTGCTCTG-3’ and *MBP* (OLIG2 BS): 5’-AAAGCAGCTGCCTGG-3’. The following primers were used for ChIP-qPCR analysis: *MBP* (MYRF ChIP): (f: 5’-GTCTGCTTGTCTGTTTCCCA-3’, r: 5’-TCCGGGGATTGACTTTCAGA-3’), *SIP1* (OLIG2 ChIP): (f: 5’-TCCTCATGGAACTTGAGTCG-3’, r: 5’-AAAACCCTGATATCCTGACAGC-3’), *CNP* (OLIG2 ChIP): (f: 5’-GAA CCTGGACCAAAGCTGAC-3’, r: 5’-TCTGAGGTGTCTCCCCACTC-3’) and *MBP* (OLIG2 ChIP): (f: 5’-TCGTGTGCAGTGCTTAGACC-3’, r: 5’-TTTCATTCAGCCTGTGCTTG-3’). The PCR was run with the following conditions: 1. Hotstart: 3 min 95°C, 2. Amplification: 30 sec 95°C, 30 sec 63°C, 30 sec 72°C (40 cycles). ChIP-qPCR was analyzed by calculating the relative amount of immunoprecipitated DNA compared to INPUT DNA for the control regions (% of recovery) using the following formula: % recovery = 2^[(Ct_input_ – 6.64) - Ct_sample_)] x 100%.

### Statistical analysis

*Prism 8* software (GraphPad Software Inc.) was used for statistical analysis. For all experimental values, mean and standard error of the mean (SEM) are given. Shapiro-Wilk normality test was applied for distribution testing. Group comparisons were performed with one-way ANOVA or Kruskal-Wallis test and Tukey’s or Dunn’s multiple comparison post hoc test. Significance thresholds were set to: *P < 0.05, **P < 0.01, ***P < 0.001. Correlations of axon diameter with myelin sheath thickness, g-ratio and mRNA expression levels with transcription factor binding capacities were tested using Pearson’s correlation. Data for group comparisons for cumulative frequencies were tested for normality and logarithmically transformed before applying the one-way ANOVA and Tukey’s multiple comparison post hoc test.

## Results

### MR imaging and histopathological analysis of FCD type II cases

All patients included in this study were scanned by MRI with either FLAIR- or STIR-weighted sequences according to a standard protocol to detect potential lesions (Wellmer et al., 2013, Urbach et al., 2015, Bernasconi et al., 2019). The FCD IIa group was heterogeneous with respect to a detectable lesion: five patients showed a thickening of the cortex and blurring of the gray-white matter boundary (Figure 1a, b), one case had a transmantle sign and three were MRI-negative (T, all patients of the FCD IIb cohort presented with a transmantle sign, a zone of increased signal intensity tapering toward the lateral ventricle (Figure 1c, d; Table 1).

**Figure 1.**
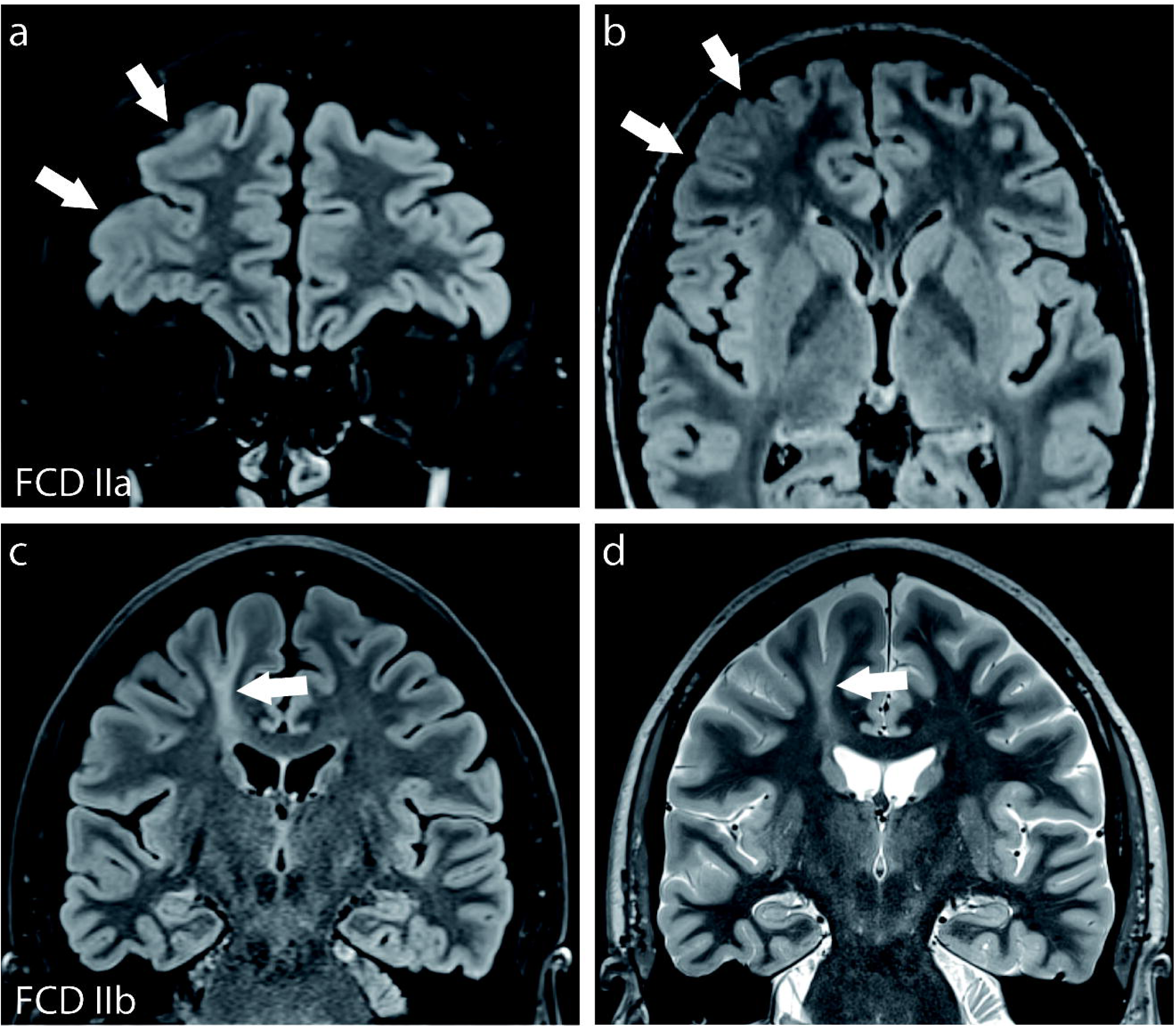
Presurgical MRI of FCD IIa and IIb patients. Representative 3T MRI of two patients with frontal FCD IIa (a, b) and FCD IIb (c, d) illustrating their differing macroscopic phenotype. Subtle cortical thickening and blurring is seen at the gray-white matter junction in FCD IIa (arrows in a, b; a: coronal, b: axial, both FLAIR). FCD IIb presents with a transmantle and deep sulcus sign (arrows in c, d; c: FLAIR, d: T2 STIR; both coronal).

Since FCDs can be variable in extensions and dimensions, we characterized every resected specimen included in this study regarding the lamination pattern and cytoarchitecture and determined the FCD type according to the ILAE classification (Blümcke et al., 2011, Najm et al., 2022). To this end, serial sections of all cases were stained with cresyl violet and were immunolabeled for NeuN to visualize all layers, or for SMI32, a monoclonal antibody specific for non-phosphorylated neurofilament H (Sternberger and Sternberger, 1983), which labels pyramidal cells of layers III and V.

In controls, all six cortical layers were present as shown by Nissl staining or NeuN labeling (Figure 2a, b). Pyramidal cells in layers III and V appeared normally shaped with parallel-orientated apical dendrites, which reached the border of layer I (Figure 2c). In the FCD IIa cohort, Nissl and NeuN stainings showed a mild hypercolumnization across layers II to V (Figure 2d, e). Moreover, the occurrence of dysmorphic neurons, characterized by malorientation and the strong accumulation of non-phosphorylated neurofilament was apparent, in particular, in layers III and V/VI (Figure 2f, j). FCD IIb cases were characterized by blurring of the cortical layers (Figure 2g, h), the presence of dysmorphic neurons in all layers and at the gray-white matter boundary (Figure 2i, k). Balloon cells, characteristic for FCD IIb, were recognizable in the cortical white matter by vimentin labeling (Figure 2l).

**Figure 2.**
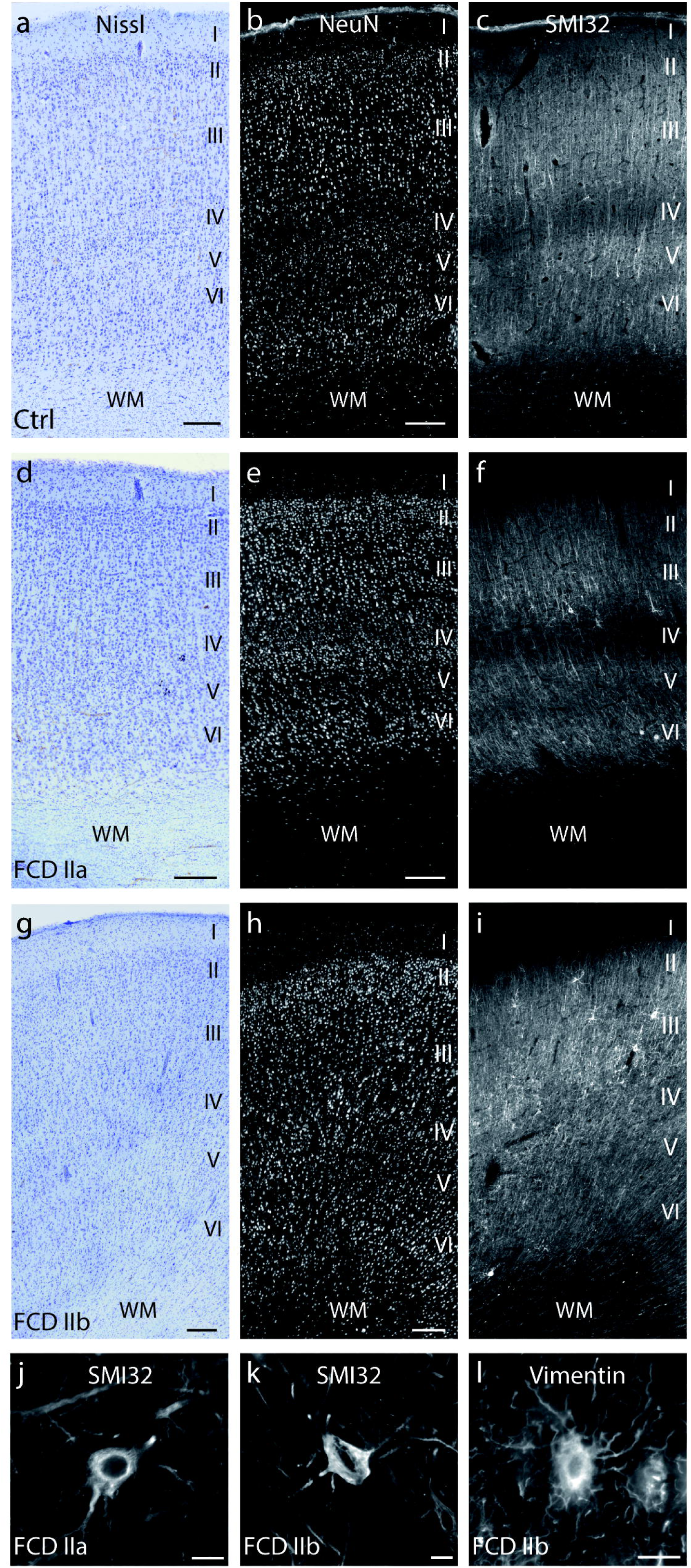
Histological characteristics of epileptic non-dysplastic control, FCD IIa and FCD IIb specimens in the frontal lobe. Representative photomicrographs of cresyl violet-stained (a, d, g) and NeuN-(all neurons) (b, e, h) or SMI32-immunolabeled (pyramidal neurons in layers III and V) tissue sections (c, f, i). The FCD IIa case shows a radial lamination disturbance with hypercolumnization (d, e). In controls and FCD IIa, SMI32-positive pyramidal neurons are densely arranged in layers III and V (c, f) with the appearance of dysmorphic neurons across all layers in FCD IIa but concentrated in supragranular layers in FCD IIb (i). Detailed images of SMI32-positive cells reveal the characteristic shape of dysmorphic neurons in FCD IIa (j) and FCD IIb (k). The hallmark of FCD IIb is the presence of Balloon cells which are visualized by vimentin staining (l). WM: white matter; Scale bars: (a-i) 250 µm; (j-l) 20 µm. I-VI: cortical layers.

Altogether, by applying presurgical MRI and histological characterization of all specimens, we ensured the characteristic phenotype of the two FCD cohorts and of the control group.

### Distribution of myelinating OLs in the gray matter of frontal lobe FCD types II

Based on our pervious results from the temporal lobe, where we had found FCD-associated myelination deficits and reduced numbers of myelinating OLs (Donkels et al., 2017, Donkels et al., 2020), we focused in the current study on frontal lobe FCD IIa and IIb. We performed ISHH for *mbp* mRNA, which is exclusively expressed in mature, myelinating OLs (Baumann and Pham-Dinh, 2001), to determine the distribution of the myelin-producing cells (Figure 3). In controls, *mbp* mRNA-expressing OLs were most numerous in the infragranular layers of the gray matter and thinned out gradually up to layer I (Figure 3b, c). In FCD IIa, a similar distribution was detectable but some OLs, located in the deep cortical layers, exhibited very strong *mbp* mRNA signals (Figure 3e, f). In contrast, in FCD IIb *mbp* mRNA-expressing OLs appeared more numerously across the entire cortex when compared to controls and FCD IIa cases (Figure 3h, i). Indeed, cell counting revealed significantly more *mbp* mRNA-expressing OLs in FCD IIb when compared to FCD IIa [controls (n=6): 186.6 ± 28.16 cells/mm²; FCD IIa (n=9): 168.3 ± 25.35 cells/mm²; FCD IIb (n=9): 276.5 ± 33.36 cells/mm^2^; (one-way ANOVA, F=4.084, p=0.0317;Tukey’s post hoc test: control vs. FCD IIa p=0.9118, control vs. FCD IIb p=0.1329, FCD IIa vs. FCD IIb p=0.0334)] (Figure 3j).

**Figure 3.**
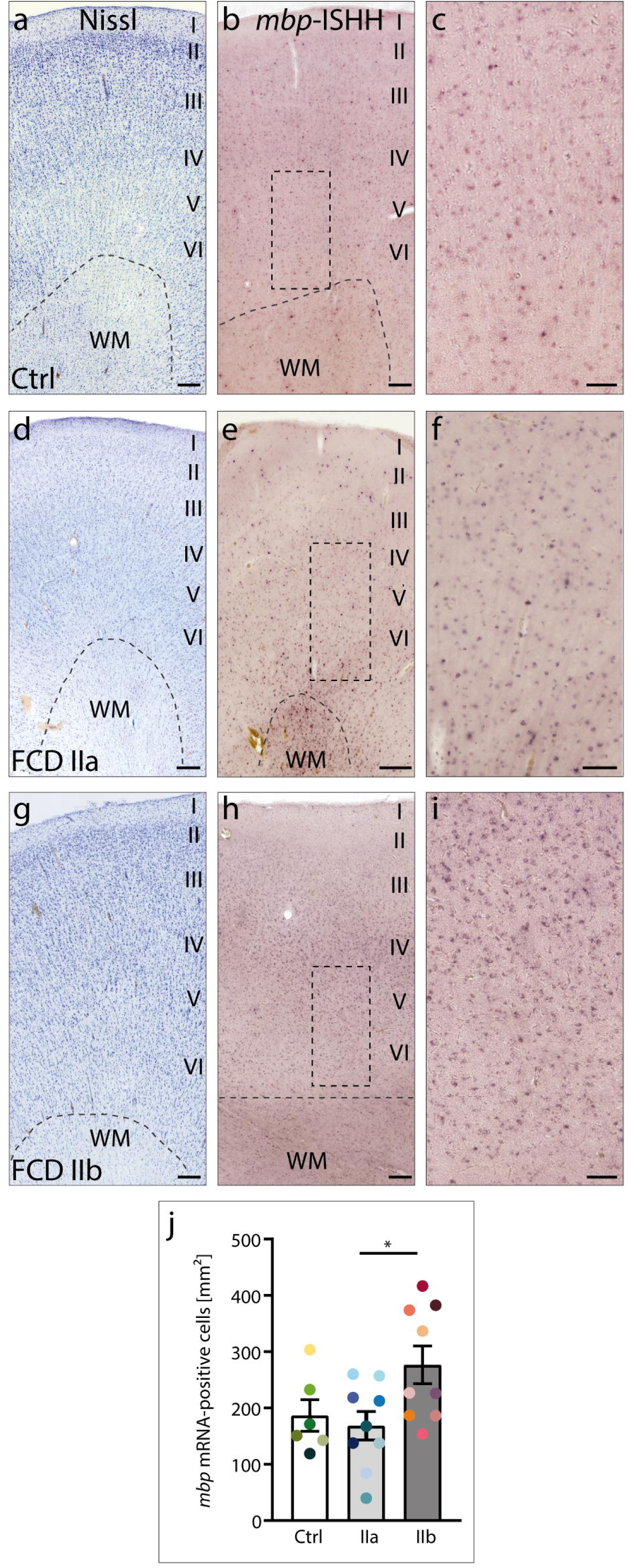
Distribution of *mbp* mRNA-expressing OLs in the gray matter of frontal lobe control and FCD type II specimens. Representative photomicrographs of cresyl violet-stained (a, d, g) and *mbp-*ISHH-processed (b, c, e, f, h, i) tissue sections of controls (a-c), FCD IIa (d-f) and FCD IIb (g-i) cases. ROIs in (b, e, h) indicate the magnified areas shown on the right. *Mbp* mRNA-expressing OLs are loosely distributed across the gray matter. Counting of these cells in all cortical layers reveals significantly higher cell numbers in FCD IIb when compared to FCD IIa (one-way ANOVA F=4.084, p=0.0317; Tukey’s post hoc test: FCD IIa vs. FCD IIb p=0.0334) (j). Individual patients are color-coded: greenish: control, blueish: FCD IIa, redish: FCD IIb. WM, white matter; scale bars: (a, b, d, e, g, h) 250 μm; (c, f, i) 100 µm, I-VI, cortical layers

Altogether, FCD IIb was characterized by increased numbers of myelinating OLs, whereas FCD IIa remained unchanged regarding cell densities in the frontal lobe.

### Distribution and architecture of myelinated fibers in the gray matter of FCD IIa and IIb

To investigate the architecture of myelinated fibers in the gray matter of control and FCD type II specimens, we used immunolabeling for CNPase, a marker for pre-OLs, mature OLs and myelin (Vogel and Thompson, 1988).

In control and FCD IIa tissue, a dense, radially aligned network of CNPase-positive myelinated fibers extended from the gray-white matter boundary up to layer III, declining gradually from the white matter to the upper layers (Figure 4a, b). This pattern was present in all FCD IIa cases. High resolution confocal microscopy of layer V showed the regularly aligned myelinated fibers and OL cell bodies in controls and FCD IIa (Figure 4e, f).

**Figure 4.**
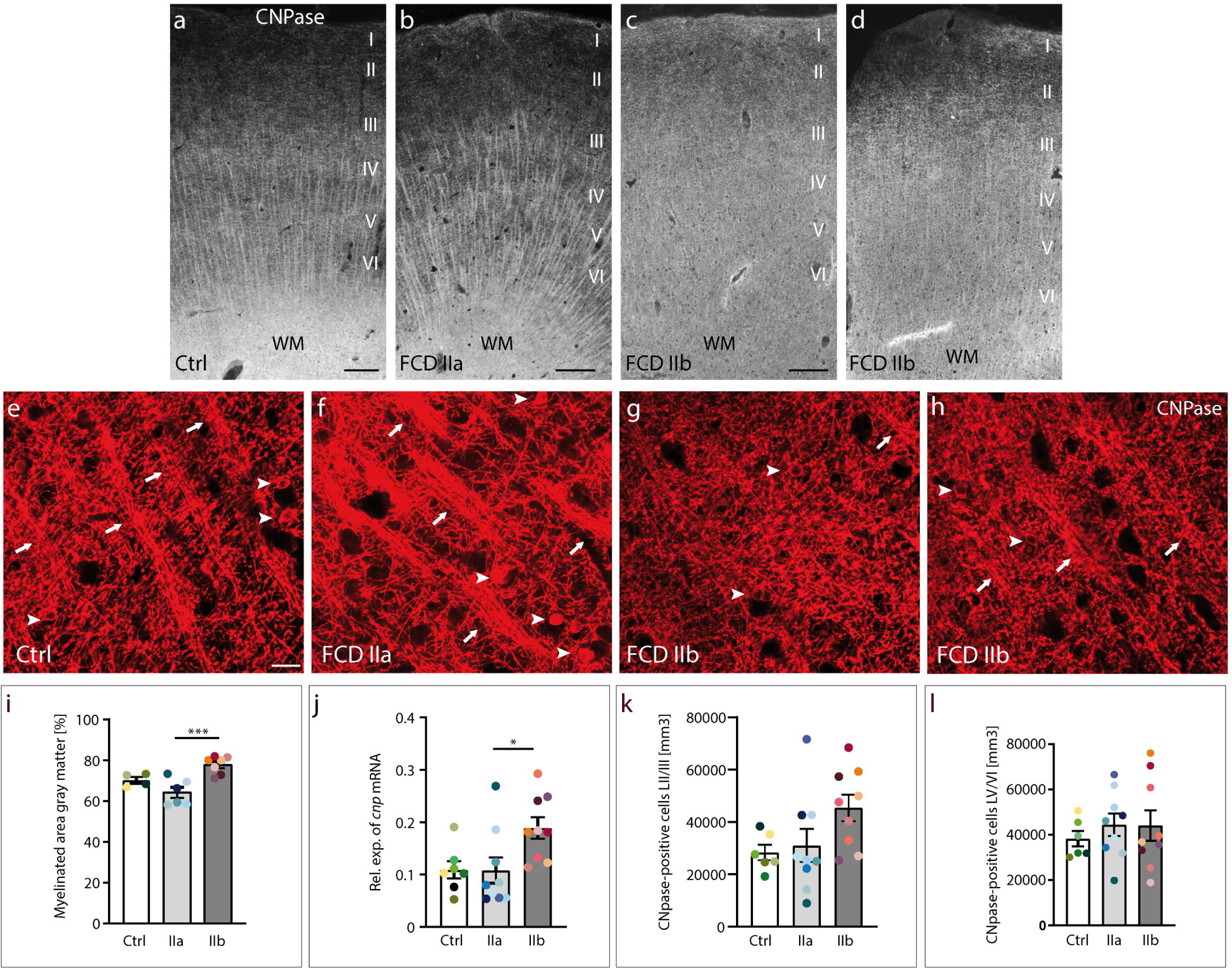
Myelination pattern in control, FCD IIa and IIb cases from the frontal lobe. Representative photomicrographs of tissue sections immunolabeled for CNPase (marker for myelin, pre-OLs and OLs) in the gray matter of control (a), FCD IIa (b) and FCD IIb (c, d) cases. The radially aligned CNPase-positive fibers reach the border of layer III in control and FCD IIa (a, b). In FCD IIb, CNPase-immunolabeled fibers reach up to layer II and show a disparate distribution with areas of disorganization (c) or radial alignment (d). High resolution confocal imaging in layers V and VI reveals the dense network of myelinated fibers and myelinating OLs and pre-Ols (e-h). Individual micrographs of confocal stacks are shown. In controls and FCD IIa, horizontal and radial CNPase-labeled fiber strands (arrows), pre-OL and OL cell bodies (arrow heads) are present (e, f). In FCD IIb, this pattern of myelinated fibers is either severely disorganized (g) or less developed (h). (i) The percentage of the myelinated area in the gray matter is significantly larger in FCD IIb compared to control and FCD IIa (one-way ANOVA, F=12.01; p=0.0009). (j) RT-qPCR demonstrates significantly increased *cnp* mRNA levels in FCD IIb compared to FCD IIa (Kruskal-Wallis p=0.0244, Dunn’s multiple comparisons test: p=0.0375). (k, l) Quantification of CNPase-expressing pre-OL and OL cell bodies in layers II/III (k) and V/VI (l) separately reveals similar cell numbers in all cases (layers II/III: one-way ANOVA, F=2.910, p=0.0766; layers V/VI: one-way ANOVA, F=0.3356, p=0.7187). Colored dots identify individual patients, greenish: control, blueish: FCD IIa, redish: FCD IIb. Scale bars: (a-d) 250 μm; (e-h) 20 µm; WM, white matter. I-VI: cortical layers.

In all FCD IIb cases, however, the radial CNPase-positive fiber pattern was much weaker and inhomogeneously distributed. In the same slice, areas with radial fiber pattern reaching up to layer II (Figure 4d) were next to regions with disorganized myelin fibers (Figure 4c). At high resolution, this irregular pattern was clearly apparent with hardly any individual strands (Figure 4g) or a marginal fiber architecture (Figure 4h). When we measured the percentage of the myelinated area in the gray matter, we found a significantly larger myelinated area in FCD IIb (n=7) compared to FCD IIa (n=6) but not to controls (n=4) (one-way ANOVA, F=12.01, p=0.0009; Tukey’s post hoc test: control vs. FCD IIa p=0.1765, control vs. FCD IIb p=0.0767, FCD IIa vs. FCD IIb p=0.0007) (Figure 4i). This was also reflected by significantly increased *cnp* mRNA levels (Kruskal-Wallis, p=0.0244, Dunn’s multiple comparisons test: control vs. FCD IIa p>0.9999, control vs. FCD IIb p=0.0992, FCD IIa vs. FCD IIb p=0.0375) (Figure 4j). However, counting of CNPase-labelled pre-OL and OL cell bodies in layers II/III and V/VI separately did not reveal any significant differences in cell numbers between controls, FCD IIa and FCD IIb cases [layers II/III: controls (n=6): 28346 ± 2935 cells/mm^3^; FCD IIa (n=9): 31062 ± 6296 cells/mm^3^; FCD IIb (n=9): 45400 ± 5016 cells/mm^3^; (one-way ANOVA, F= 2.910, p=0.0766)] (Figure 4k); [layers V/VI: controls: 38207 ± 3422 cells/mm^3^; FCD IIa: 44408 ± 4957 cells/mm^3^; FCD IIb: 44268 ± 6714 cells/mm^3^; (one-way ANOVA, F= 0.3206, p=0.7292)] (Figure 4l).

In summary, this analysis revealed an enlarged myelinated area, an irregular myelination pattern and increased *cnp* mRNA expression levels only in the gray matter of FCD IIb but not in FCD IIa.

### Ultrastructure of myelinated axons in the gray matter of frontal lobe FCD types II

Next, we compared the ultrastructure of myelinated axons of control and FCD II cases by transmission electron microscopy in layers V/VI (Figure 5a-c), where axons show a high and homogeneous myelin coverage (Tomassy et al., 2014). We determined the axon diameter, the myelin sheath thickness and the g-ratio, defined as the ratio of the axon diameter to the total fiber diameter (Rushton, 1951).

**Figure 5.**
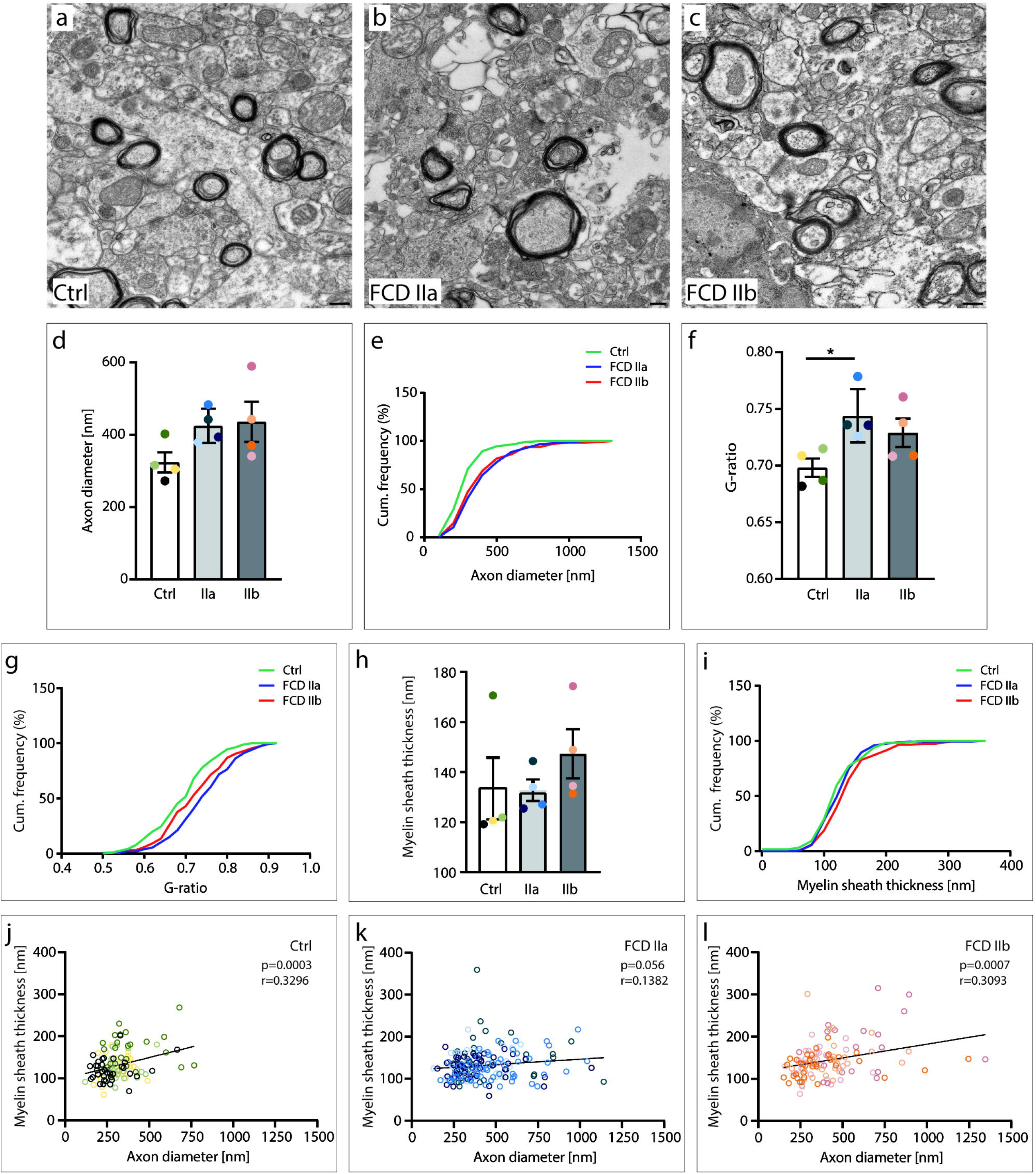
Ultrastructure of myelinated axons in FCD II types of the frontal lobe. Representative electron photomicrographs of myelinated axons in layers V/VI of control (a), FCD IIa (b) and FCD IIb (c) specimens. (d) Bar graph demonstrating that the mean axon diameters of FCD IIa and FCD IIb have on average larger axon diameters than control cases (one-way ANOVA, F=2.59, p=0.129). (e) The cumulative frequency distribution of axon diameters shows significantly increased axon diameters in both FCD II types (one-way ANOVA, F=19.26, p<0.0001; Tukey’s post hoc test: control vs. FCD IIa p<0.0001, control vs. FCD IIb p<0.0001). (f) The mean g-ratio value is significantly increased in FCD IIa cases when compared to controls (one-way ANOVA, F=4.511, p=0.044; Tukey’s post hoc test: control vs. FCD IIa p=0.039). (g) The cumulative frequency distribution of g-ratios shows significant higher g-ratios for FCD IIb and FCD IIa (one-way ANOVA, F=14.69, p<0.0001; Tukey’s post hoc test: control vs. FCD IIa p<0.0001, control vs. FCD IIb p=0.019, FCD IIa vs. FCD IIb p=0.046). (h) Bar graph demonstrating the mean myelin sheath thickness which is similar in the three analyzed groups (one-way ANOVA F=0.746, p=0.501). (i) The cumulative frequency distribution of myelin sheath thickness does not reveal differences between controls and FCD IIa but a significant increase of FCD IIb in comparison to FCD IIa and controls (one-way ANOVA, F=4.487, p=0.012; Tukey’s post hoc tests: control vs. FCD IIb p=0.023, FCD IIa vs. FCD IIb p=0.024). (j-l) Correlation plots of axon diameters versus myelin sheath thickness. (j) Controls, (k) FCD IIa and (l) FCD IIb. There is a significant correlation of axon diameters and the thickness of myelin sheaths for controls (Pearson’s correlation: r=0.329, p=0.0003) and FCD IIb (Pearson’s correlation: r=0.309, p=0.0007) but not for FCD IIa (Pearson’s correlation: r=0.138, p=0.056). Colored dots represent individual patients, greenish: control, blueish: FCD IIa, redish: FCD IIb. Scale bars (a-c): 250 nm.

Mean axon diameters were slightly increased in both FCD cohorts when compared to the control group [(axon diameters: control: 323.9 ± 27.7 nm; FCD IIa: 424.8 ± 23.8 nm; FCD IIb: 436.1 ± 55.5 nm); (one-way ANOVA F=2.59, p=0.129)] (Figure 5d). When we looked at the frequency distribution of all axon diameters within each group, we found a significant shift towards larger axon diameters in both FCD types [controls (n=4; 114 axons) 119.9.8 – 767.5 nm; FCD IIa (n=4; 192 axons) 129.5 – 1141.5 nm; FCD IIb (n=4; 116 axons) 151.4 – 1346.2 nm] (one-way ANOVA F=19.26, p<0.0001; Tukey’s post hoc test: control vs. FCD IIa p<0.0001, control vs. FCD IIb p<0.0001, FCD IIa vs. FCD IIb p=0.691)] (Figure 5e). The mean g-ratio of myelinated axons was elevated in FCD IIb and significantly increased in FCD IIa when compared to non-dysplastic controls [(g-ratios: control: 0.698 ± 0.008; FCD IIa: 0.744 ± 0.017; FCD IIb: 0.728 ± 0.012); (one-way ANOVA F=4.511, p=0.044; Tukey’s post hoc test: control vs. FCD IIa p=0.039, control vs. FCD IIb p=0.174, FCD IIa vs. FCD IIb p=0.611)] (Figure 5f). This was even more prominent when we looked at the relative frequency distribution of all g-ratio values: both FCD II groups had significantly increased g-ratios in comparison to the control group and FCD IIa showed significantly higher values than FCD IIb [(one-way ANOVA F=14.69, p<0.0001; Tukey’s post hoc test: control vs. FCD IIa p<0.0001, control vs. FCD IIb p=0.019, FCD IIa vs. FCD IIb p=0.046)] (Figure 5g). Normally, an increased g-ratio indicates a reduced thickness of the myelin sheath, based on the assumption that the axon diameters are not altered. However, since our FCD cases had significantly larger axon diameters, we investigated also the myelin sheath thicknesses. On average, there was no difference between the three groups [(myelin sheath thickness: control: 133.5 ± 12.39 nm; FCD IIa: 132.7 ± 4.312 nm; FCD IIb: 147.3 ± 9.828 nm); (one-way ANOVA F=0.746, p=0.501)] (Figure 5h). However, plotting the relative frequency distribution of all individual values revealed that FCD IIb had significantly thicker myelin sheaths when compared to controls and FCD IIa cases [(one-way ANOVA F=4.487, p=0.012; Tukey’s post hoc test: control vs. FCD IIa p= 0.941, control vs. FCD IIb p=0.023, FCD IIa vs. FCD IIb p=0.024)] (Figure 5i) but also thicker axons than controls.

So solve this controversy, we correlated all individual axon diameters with the corresponding myelin sheath thickness for the three groups separately: (in nm): controls (range: smallest axon diameter/myelin sheath 119.92/91.59; largest axon diameter/myelin sheath 767.5/130.77); FCD IIb (range: smallest axon diameter/myelin sheath 151.41/103.22, largest axon diameter/myelin 1346.19/146.09). We found significant correlations between axon diameter and myelin sheath thickness in controls (Pearson’s correlation: r=0.329, p=0.0003) and FCD IIb (Pearson’s correlation: r=0.309, p=0.0007) (Figure 5j, l), implying that the axon caliber determines the thickness of the myelin sheath (Stassart et al., 2018). This was not the case for FCD IIa (Pearson’s correlation: r=0.138, p=0.056) (range: smallest axon diameter/myelin sheath 129.48/124.41, largest axon diameter/myelin sheath 1141.5/92.95), where large caliber axons (above 700 nm) had thinner myelin sheaths than axons of the same caliber in controls or FCD IIb (Figure 5k) indicating that myelination was selectively compromised for axons with a large diameter in FCD IIa.

### Transcriptional regulation of OL differentiation factors and myelin-associated genes

Since thinner myelin sheaths may indicate a disturbance in OL differentiation and/or myelin synthesis, we next tested whether the transcriptional regulation of OL differentiation and myelination was affected in our FCD cohorts. First, we performed real-time RT-qPCR to measure the mRNA levels of transcription factors either sequentially active at different stages of OL maturation (SIP1, MYRF) or continuously expressed (OLIG2) (Emery and Lu, 2015). In addition, we quantified the expression levels of *mbp*, *mag* and *mog* mRNAs, which encode proteins that are all components of the myelin sheath. *Olig2* (Kruskal-Wallis, p=0.6391) and *sip1* (one-way ANOVA, F=1.487, p=0.2480) mRNA levels were similar in controls (n=7), FCD IIa (n=9) and FCD IIb (n=9) (Figure 6a, b), whereas *myrf* mRNA expression was significantly up-regulated in FCD IIb when compared to controls and FCD IIa (one-way ANOVA, F=9.054, p=0.0014; Tukey’s post hoc test: control vs. FCD IIa p=0.7367, control vs. FCD IIb p=0.0159, FCD IIa vs. FCD IIb p=0.0015) (Figure 6c). Likewise, *mag* (one-way ANOVA, F=12.64, p=0.0002; Tukey’s post hoc test: control vs. FCD IIa p=0.8064, control vs. FCD IIb p=0.0029, FCD IIa vs. FCD IIb p=0.0003), and *mog* (one-way ANOVA, F=7.302, p=0.0037; Tukey’s post hoc test: control vs. FCD IIa p=0.7709, control vs. FCD IIb p=0.0323, FCD IIa vs. FCD IIb p=0.0039), mRNA levels were also significantly increased in FCD IIb in comparison to controls and FCD IIa (Figure 6e, f) reflecting the regulatory action of MYRF on myelin genes. Similarily, *mbp* mRNA levels were increased in FCD IIb when compared to the FCD IIa cohort, however, no significant difference was found between FCD IIb and controls (Kruskal-Wallis, p= 0.0016; Dunn’s multiple comparisons test: control vs. FCD IIa p=0.2230, control vs. FCD IIb p=0.3488, FCD IIa vs. FCD IIb p=0.0010). These results point to an FCD-associated transcriptional dysregulation in the late OL differentiation stage.

**Figure 6.**
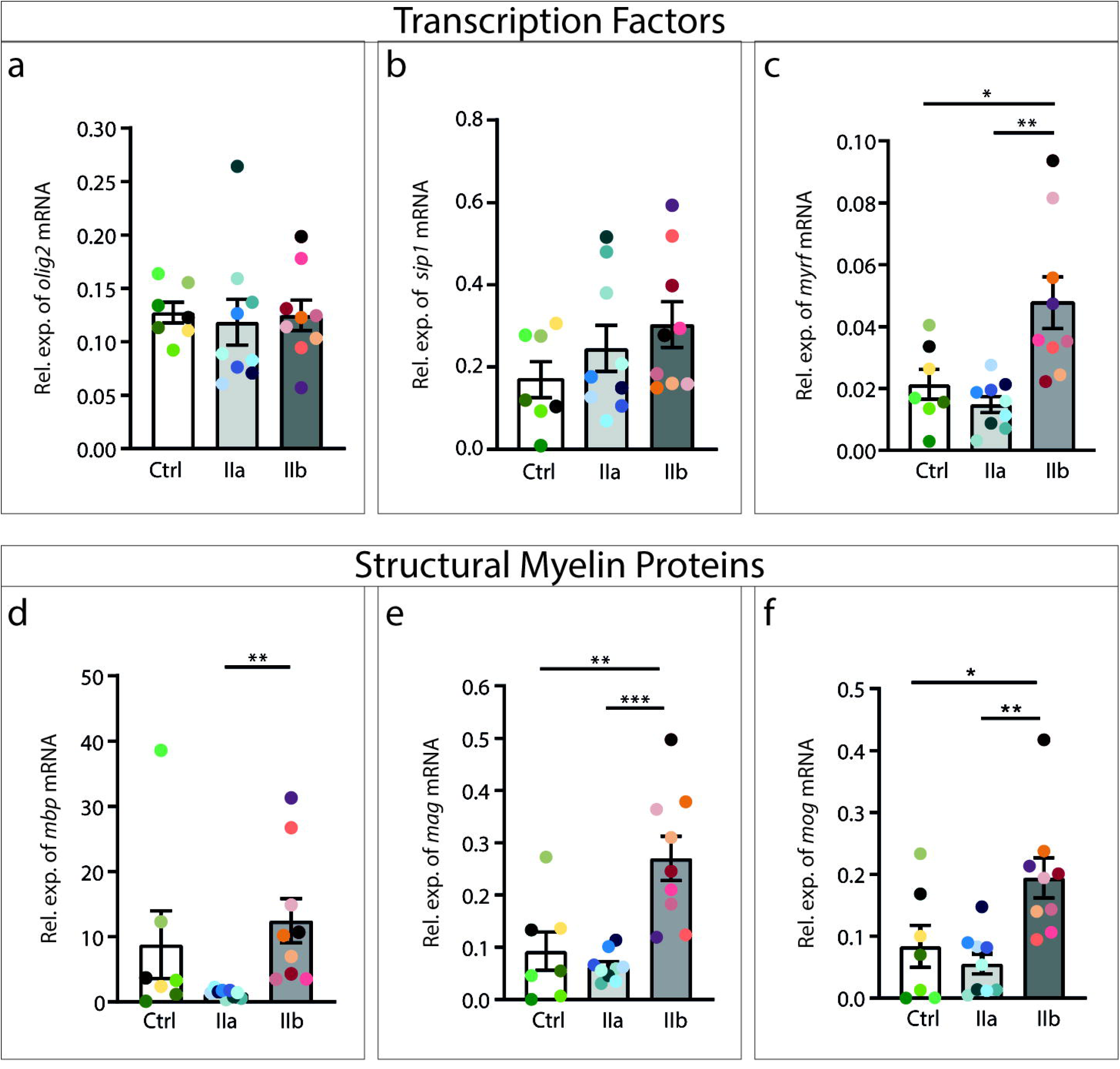
Expression levels of transcription factor and myelin-associated transcripts in frontal lobe FCD IIa and IIb. All mRNA levels were quantified by real-time RT-qPCR. The expression of the transcription factor mRNAs *olig2* (Kruskal-Wallis p=0.6391) (a) and *sip1* (one-way ANOVA, F=1.487, p=0.2480) (b) are similar in all groups. The mRNA levels of *myrf* (c) and of the myelin components *mag* (e) and *mog* (f) are significantly increased in FCD IIb when compared to controls and reduced in FCD IIa when compared to FCD IIb [(*myrf*: one-way ANOVA, F=9.054, p=0.0014; Tukey’s post hoc test: control vs. FCD IIa p=0.7367, control vs. FCD IIb p=0.0159, FCD IIa vs. FCD IIb p=0.0015), (*mag*: one-way ANOVA F=12.64, p=0.0002; Tukey’s post hoc test: control vs. FCD IIa p=0.8064, control vs. FCD IIb p=0.0029, FCD IIa vs. FCD IIb p=0.0003), (*mog*: one-way ANOVA F=7.302, p=0.0037; Tukey’s post hoc test: control vs. FCD IIa p=0.7709, control vs. FCD IIb p=0.0323, FCD IIa vs. FCD IIb p=0.0039)]. There is a significant increase of *mbp* mRNA level in FCD IIb when compared to FCD IIa (*mbp*: Kruskal-Wallis, p=0.0016; Dunn’s multiple comparisons test: control vs. FCD IIa p=0.2230, control vs. FCD IIb p=0.3488, FCD IIa vs. FCD IIb p=0.0010). Colored dots represent individual patients, greenish: control, blueish: FCD IIa, redish: FCD IIb.

To target transcriptional regulation directly, we next performed ChIP assays which give insight into transcription factor binding to specific promoters. Chromatin was purified from gray matter of control, FCD IIa and IIb specimens and immunoprecipitated with antibodies specific for transcription factors which either determine the entire OL lineage (OLIG2) or the advanced OL differentiation stage (MYRF). As transcription factor targets, we chose either genes encoding transcription factors (*SIP1, SOX10*) or structural proteins of the myelin sheath (*MBP*, *MAG*, *MOG*, *CNP*) and quantified the OLIG2 and MYRF binding capacities to the predicted binding sites by real-time qPCR.

We did not find any differences in the binding capacities of OLIG2 to the *SIP1* (one-way ANOVA F=0.3313, p=0.7215), *MBP* (one-way ANOVA F=0.2883, p=0.7525), *CNP* (one-way ANOVA F=0.6954, p=0.5100) and *MOG* (Kruskal-Wallis p=0.9582) promoters between control (n=7), FCD IIa (n=9) and FCD IIb (n=9) cases indicating that OLIG2 is equally active in the different groups (Figure 7a-d). In contrast, MYRF showed a significantly decreased affinity to the *SOX10* promotor in FCD IIa when compared to the control group (one-way ANOVA F=5.006, p=0.0162; Tukey’s post hoc test: control vs. FCD IIa p=0.0228, control vs. FCD IIb p=0.8562, FCD IIa vs. FCD IIb p=0.0505) (Figure 7e). Additionally, binding capacities of MYRF to the *MBP* (one-way ANOVA F=4.965, p=0.0166; Tukey’s post hoc test: control vs. FCD IIa p=0.1541, control vs. FCD IIb p=0.6000, FCD IIa vs. FCD IIb p=0.0137) and *MAG* promoters (Kruskal-Wallis p=0.0431; Dunn‘s multiple comparison test: control vs. FCD IIa p=0.2828, control vs. FCD IIb p>0.999, FCD IIa vs. FCD IIb p=0.0448) were increased in FCD IIb specimens when compared to FCD IIa (Figure 7f, g). Altogether, these results indicate an FCD-associated transcriptional dysregulation of myelin-associated genes in the final OL differentiation stage.

**Figure 7.**
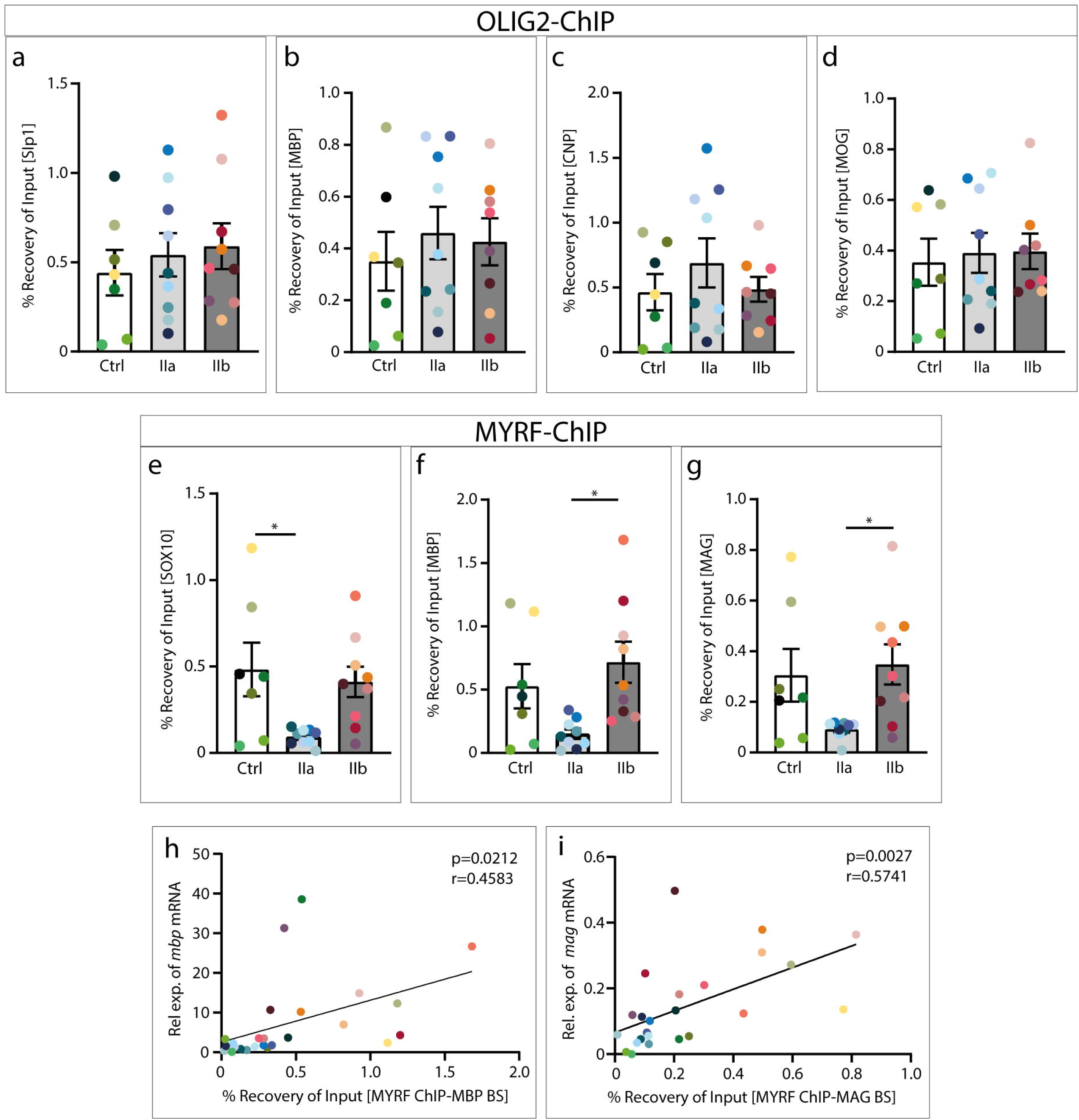
Transcription factor binding capacities to promoters of myelin-associated genes. *OLIG2* (a-d) and *MYRF* (e-g) binding sites were precipitated using ChIP assays and quantified using real-time qPCR. The amounts of precipitated DNA are presented as the percentage of input DNA. The binding capacities of OLIG2 to the *SIP1* (a), *MBP* (b), *CNP* (c) and *MOG* (d) promoters are similar in all analyzed groups [(OLIG2 on *SIP1*: one-way ANOVA, F=0.3313, p=0.7215), (OLIG2 on *MBP*: one-way ANOVA F=0.2883, p=0.7525), (OLIG2 on *CNP*: one-way ANOVA, F=0.6954, p=0.5100), (OLIG2 on *MOG*: Kruskal-Wallis p=0.9582)]. However, MYRF has significantly lower binding potential to the *SOX10* promotor (e) in FCD IIa when compared to control (MYRF on *SOX10*: one-way ANOVA, F=5.006, p=0.0162; Tukey’s post hoc test: control vs. FCD IIa p=0.0228, control vs. FCD IIb p= 0.8562, FCD IIa vs. FCD IIb p= 0.0505) and to the *MBP* (f), and *MAG* (g) promoters in FCD IIa when compared to FCD IIb [(MYRF on *MBP:* one-way ANOVA F=4.965, p=0.0166; Tukey’s post hoc test: control vs. FCD IIa p=0.1541, control vs. FCD IIb p=0.6000, FCD IIa vs. FCD IIb p=0.0137), (MYRF on *MAG:* Kruskal-Wallis p=0.0431; Dunn‘s multiple comparison test: control vs. FCD IIa p=0.2828, control vs. FCD IIb p>0.9999, FCD IIa vs. FCD IIb p=0.0448)]. Pearson’s correlation of MYRF binding capacities with *mpb* (i) and *mag* (j) mRNA expression levels was performed for the percentage of input values from the ChIP assays with the relative expression values of the RT-qPCRs. Significant correlations are present between MYRF binding capacities to *MBP* and *MAG* promoters and *mpb* (i) and *mag* (j) mRNA levels, in particular for FCD IIb (Pearson’s correlations for *mbp*: r=0.4583, p=0.0212 and *mag*: r=0.5741, p=0.0027). Colored dots represent individual patients, greenish: control, blueish: FCD IIa, redish: FCD IIb.

Finally, we tested, whether there is a direct impact of the investigated transcription factor binding capacities on the corresponding mRNA expressions. We correlated the promoter binding capacities of OLIG2 and MYRF with the corresponding *sip1*, *mbp*, *mog*, and *mag* mRNA levels. There were no differences in the promoter binding capacities of OLIG2 to their target promoters (data not shown). However, we found a significant correlation of the MYRF binding capacity with expression levels of *mbp* (Pearson’s correlation: r=0.4583, p=0.0212) and *mag* transcripts (Pearson’s correlation: r=0.5741, p=0.0027) in FCD IIb but less in FCD IIa (Figure 7h, i) mirroring the high expression of myelin-associated genes in FCD IIb.

## Discussion

Insulation of axons by myelin is important for fast and long-range signal propagation in the brain. Myelin is constantly produced in the adult brain by oligodendrocytes in a plasticity-dependent manner and necessary for memory consolidation (Steadman et al., 2020). Myelination deficiencies have been found in several neurologic disorders in particular in multiple sclerosis (Nave and Werner, 2014) and are often associated with seizure occurrence (Rayatpour et al., 2021). There is increasing awareness that also in epilepsy maladaptive myelination or axon demyelination are of high relevance for seizure generation (De Curtis et al., 2021, Knowles et al., 2022).

In the current study, we investigated maladaptive myelination and the transcriptional regulation of myelin-associated genes in FCD II specimens obtained from patients with pharmacoresistant focal epilepsy. We focused on the gray matter of FCD IIa and IIb cases from the human frontal lobe, all thoroughly characterized by presurgical MR scans and *post hoc* histological examination. In the MRT, most FCD IIa patients showed a thickening of the neocortex and blurring of the gray-white matter boundary, whereas all FCD IIb cases displayed a transmantle sign (Urbach et al., 2002). Histological characterization ensured that all specimens of our study fulfilled the morphological criteria of FCD IIa and IIb proposed by the ILAE (Najm et al., 2022).

The myelinated area and extension of radially orientated, myelinated CNPase-positive fibers, was similar in FCD IIa and controls, reaching up to layer III as described earlier for the temporal cortex (Zucca et al., 2016), (Donkels et al., 2020). Likewise, the structural organization of myelinated fiber strands was intact in FCD IIa. This is in contrast to the temporal lobe, where CNPase-positive fibers have been shown to be strongly disorganized in FCD IIa (Donkels et al., 2017) pointing to a brain region-dependent phenotype. FCD IIb cases, however, presented with an enlarged myelinated area characterized either by a completely disorganized fiber pattern or partial maintenance of radially orientated fibers extending up to layer II as shown by epifluorescence and confocal laser microscopy. A similar disorganization of myelinated fibers has been described for extra-temporal FCD IIb by Shepherd et al. (2013) suggesting an FCD IIb-specific myelination disturbance.

When we analyzed myelinated axons ultrastructurally and determined axon diameter, myelin sheath thickness and g-ratio, we found a significantly increased mean g-ratio for FCD IIa cases, indicating a reduced thickness of the myelin sheath as found in the temporal lobe (Donkels et al., 2020). However, the frequency distribution of axon diameters indicated a significant increase of axon calibers in both FCD cohorts in comparison to controls. An observation also described by Shepherd et al. (2013) for the white matter of FCD IIb. Unfortunately, there is hardly any information about the physiological range of axon diameters in the human gray matter, since most studies focused on fiber tracts in the white matter. Liewald et al. (2014) reported a frequency distribution of 300 – 1500 nm for axon diameters in the human white matter. We found much thinner axons (119 – 767 nm) in controls but both FCD cohorts approached values (FCD IIa: 130 – 1142 nm; FCD IIb: 151 – 1346 nm) similar to those reported by Liewald et al. (2014). Since the axon diameter depends strongly on the soma size of the parent neuron (Tomasi et al., 2012), it is tempting to speculate that the presence of large dysmorphic neurons, being part of the FCD II pathology, might give rise to thicker axons in our FCD cases in comparison to control tissue.

Because the axon diameter is critical for the calculation of the g-ratio (Rushton, 1951), we performed a more refined analysis of all individual values and correlated the corresponding axon diameters and myelin sheath thicknesses. We found a significant correlation of both parameters in FCD IIb and controls, showing a linear relationship between axon diameter and myelin sheath thickness as reported in the literature (Stassart et al., 2018). This was not the case in FCD IIa specimens, where large diameter axons (≥ 700 nm) had thinner myelin sheaths than corresponding axons of FCD IIb and controls indicating a myelination deficiency of large diameter axons only in the FCD IIa cohort.

What could be the explanation for the disorganized myelin fiber structure in FCD IIb and thinner myelin sheaths in FCD IIa? It might be that the differentiation of OLs, the myelin sheath-producing glial cells, and their myelin-synthesizing capacities are compromised. In fact, a loss of myelin (Mühlebner et al., 2012), of OPCs and OLs (Scholl et al., 2017) and decreased myelin synthesis, shown in an *in vitro* myelination assay (Gruber et al., 2021), has been demonstrated for the white matter of FCD II cases. Similarly, we had found significantly less OPCs and a reduced proliferation capacity of OPCs purified from the gray matter of temporal FCD IIa (Donkels et al., 2020).

In the gray matter of the frontal lobe, investigated in the current study, the OL lineage was differentially affected in both FCD II types: we did not find altered numbers of *mbp* mRNA-expressing OLs or CNPase-positive pre-OLs and OLs in FCD IIa but higher densities of these cells in FCD IIb. Therefore, we conclude that the OL lineage is basically preserved in FCD IIa. Since FCD IIa was characterized by thinner myelin sheaths of large diameter axon, it is tempting to speculate that in FCD IIa, the mature OLs have a reduced myelination capacity as shown for white matter OLs (Gruber et al., 2021). In contrast, FCD IIb presented with increased numbers of mature, myelinating OLs accompanied by irregularities in myelinated areas and fiber architecture. As for FCD IIa, the mature OLs may be compromised in myelin sheath production and hence increase their numbers in a compensatory attempt.

This assumption is supported by an FCD IIb-specific increase of *myrf* mRNA levels, required for the differentiation and maintenance of the myelinating OL phenotype (Hornig et al., 2013). The levels of *olig2* and *sip1* mRNAs were unaltered in both FCD types in line with a constant need of these transcription factors for the maturation and preservation of OLs (Zhou and Anderson, 2002, Lu et al., 2002) and by unaltered binding capacities of OLIG2 to genes encoding structural myelin proteins as shown in our ChIP assays.

In comparison to FCD IIa and controls, FCD IIb was further characterized by enhanced expression of *mbp, mag* and *mog* mRNAs in agreement with an elevated need for myelin proteins due to thicker axons and myelin sheaths. This was also reflected on the level of transcriptional regulation analyzed by our ChIP assays: the promotor-binding capacities of MYRF to the *MBP* and *MAG* promoters were significantly elevated in FCD IIb when compared to FCD IIa indicating an enhanced myelin production. This was further supported by the significant correlation between MYRF promotor binding capacities and *mag* and *mbp* mRNA levels. Altogether, our results point to divergent forms of maladaptive myelination in FCD IIa and IIb.

What are the functional effects of an impaired myelination? Under physiological conditions, myelination depends strongly on neuronal activity (Almeida and Lyons, 2017). In rodents, there is recent evidence that neuronal activity increases OL differentiation and myelin sheath thickness in a pathway-specific manner (Gibson et al., 2014). Accordingly, Knowles et al., 2022 found increased oligodendrogenesis and myelination in an animal model for absence seizures and demonstrated that this contributes to epilepsy progression. An impact of electrical activity in human FCD type II it has been reported with myelin-associated transcripts being upregulated in electrocorticographic high-spiking areas (Srivastava et al., 2021). In contrast, we found in our human FCD IIa cases thinner myelin sheaths in layer V/VI of the temporal lobe (Donkels et al., 2020) and, as shown here, of the frontal lobe. We did not find any correlation with seizure frequency or duration (data not shown) indicating that different epilepsy types result in different forms of maladaptive myelination.

Our study demonstrates for the first time that there is a dysregulation of myelination in the gray matter of frontal lobe FCD IIa and FCD IIb. Both FCD types show divergent signs of maladaptive myelination: FCD IIa is characterized by a regular myelin fiber pattern but thinner myelin sheaths of large diameter axons and an attenuation of the myelin synthesis machinery. In contrast, FCD IIb presents a disorganized myelin fiber pattern covering a larger area, with thicker axon diameters and myelin sheaths and an elevated transcriptional turnover of myelin-associated genes. This imbalance in myelination may disrupt the equilibrium of signal conduction and thereby contribute to the epileptic phenotype.

## Authors’ contributions

CD: Conceptualization; data curation; formal analysis; project administration; visualization; writing - original draft; funding acquisition; SH: Visualization; formal analysis; CS, MJS: Neurosurgical brain resection; TD, MH, ASB, HU, MP, JB: Clinical data; UH: Statistical analysis; commenting manuscript; AV: Electron microscopy; JMN: Brain resection, clinical data from patients; CAH: Conceptualization; data interpretation, manuscript writing, resources.

## Acknowledgements

We thank all the patients who donated tissue for this study. We are grateful to Sigrun Nestel for excellent technical assistance and to Dr. Maria Stella Carro for help with establishing the ChIP method. We thank the German Research Foundation (grant number: DO 2542/1-1) and the Research Commission, Medical Faculty - University of Freiburg (grant DON1207/19) for financial support.

## References

Abdijadid S, Mathern GW, Levine MS, Cepeda C. 2015. Basic Mechanisms of Epileptogenesis in Pediatric Cortical Dysplasia. CNS Neurosci Ther 21:92–103.

Almeida RG, Lyons DA. 2017. On Myelinated Axon Plasticity and Neuronal Circuit Formation and Function. J Neurosci 37:10023–10034.

Baldassari S, Ribierre T, Marsan E, Adle-Biassette H, Ferrand-Sorbets S, Bulteau C, Dorison N, Fohlen M, Polivka M, Weckhuysen S, Dorfmüller G, Chipaux M, Baulac S. 2019. Dissecting the genetic basis of focal cortical dysplasia: a large cohort study. Acta Neuropathol 138:885– 900.

Baumann N, Pham-Dinh D. 2001. Biology of Oligodendrocyte and Myelin in the Mammalian Central Nervous System. Physiological Reviews 81:871–927.

Bernasconi A, Cendes F, Theodore WH, Gill RS, Koepp MJ, Hogan RE, Jackson GD, Federico P, Labate A, Vaudano AE, Blümcke I, Ryvlin P, Bernasconi N. 2019. Recommendations for the use of structural magnetic resonance imaging in the care of patients with epilepsy: A consensus report from the International League Against Epilepsy Neuroimaging Task Force. Epilepsia 60:1054–1068.

Blümcke I, Thom M, Aronica E, Armstrong DD, Vinters HV, Palmini A, Jacques TS, Avanzini G, Barkovich AJ, Battaglia G, Becker A, Cepeda C, Cendes F, Colombo N, Crino P, Cross JH, Delalande O, Dubeau F, Duncan J, Guerrini R, Kahane P, Mathern G, Najm I, Özkara Ç, Raybaud C, Represa A, Roper SN, Salamon N, Schulze-Bonhage A, Tassi L, Vezzani A, Spreafico R. 2011. The clinicopathologic spectrum of focal cortical dysplasias: A consensus classification proposed by an ad hoc Task Force of the ILAE Diagnostic Methods Commission1: The ILAE Classification System of FCD. Epilepsia 52:158–174.

Bujalka H, Koenning M, Jackson S, Perreau VM, Pope B, Hay CM, Mitew S, Hill AF, Lu QR, Wegner M, Srinivasan R, Svaren J, Willingham M, Barres BA, Emery B. 2013. MYRF Is a Membrane-Associated Transcription Factor That Autoproteolytically Cleaves to Directly Activate Myelin Genes. PLoS Biol 11:e1001625.

Chung C, Yang X, Bae T, Vong KI, Mittal S, Donkels C, Westley Phillips H, Li Z, Marsh APL, Breuss MW, Ball LL, Garcia CAB, George RD, Gu J, Xu M, Barrows C, James KN, Stanley V, Nidhiry AS, Khoury S, Howe G, Riley E, Xu X, Copeland B, Wang Y, Kim SH, Kang H-C, Schulze-Bonhage A, Haas CA, Urbach H, Prinz M, Limbrick DD, Gurnett CA, Smyth MD, Sattar S, Nespeca M, Gonda DD, Imai K, Takahashi Y, Chen H-H, Tsai J-W, Conti V, Guerrini R, Devinsky O, Silva WA, Machado HR, Mathern GW, Abyzov A, Baldassari S, Baulac S, Focal Cortical Dysplasia Neurogenetics Consortium, Gleeson JG, Jones M, Masser-Frye D, Sattar S, Nespeca M, Gonda DD, Imai K, Takahashi Y, Chen H-H, Tsai J-W, Conti V, Guerrini R, Devinsky O, Machado HR, Garcia CAB, Silva WA, Kim SH, Kang H-C, Alanay Y, Kapoor S, Haas CA, Ramantani G, Feuerstein T, Blumcke I, Busch R, Ying Z, Biloshytsky V, Kostiuk K, Pedachenko E, Mathern GW, Gurnett CA, Smyth MD, Helbig I, Kennedy BC, Liu J, Chan F, Krueger D, Frye R, Wilfong A, Adelson D, Gaillard W, Oluigbo C, Anderson A, Brain Somatic Mosaicism Network, Lee A, Huang AY, D’Gama A, et al. 2023. Comprehensive multi-omic profiling of somatic mutations in malformations of cortical development. Nat Genet 55:209–220.

De Curtis M, Garbelli R, Uva L. 2021. A hypothesis for the role of axon demyelination in seizure generation. Epilepsia 62:583–595.

Donkels C, Peters M, Fariña Núñez MT, Nakagawa JM, Kirsch M, Vlachos A, Scheiwe C, Schulze-Bonhage A, Prinz M, Beck J, Haas CA. 2020. Oligodendrocyte lineage and myelination are compromised in the gray matter of focal cortical dysplasia type IIa. Epilepsia 61:171–184.

Donkels C, Pfeifer D, Janz P, Huber S, Nakagawa J, Prinz M, Schulze-Bonhage A, Weyerbrock A, Zentner J, Haas CA. 2016. Whole Transcriptome Screening Reveals Myelination Deficits in Dysplastic Human Temporal Neocortex. Cereb Cortex bhv346.

Emery B. 2010. Regulation of Oligodendrocyte Differentiation and Myelination. Science 330:779–782.

Emery B, Agalliu D, Cahoy JD, Watkins TA, Dugas JC, Mulinyawe SB, Ibrahim A, Ligon KL, Rowitch DH, Barres BA. 2009. Myelin Gene Regulatory Factor Is a Critical Transcriptional Regulator Required for CNS Myelination. Cell 138:172–185.

Emery B, Lu QR. 2015. Transcriptional and Epigenetic Regulation of Oligodendrocyte Development and Myelination in the Central Nervous System. Cold Spring Harb Perspect Biol 7:a020461.

Fauser S. 2006. Clinical characteristics in focal cortical dysplasia: a retrospective evaluation in a series of 120 patients. Brain 129:1907– 1916.

Fauser S, Schulze-Bonhage A, Zentner J. 2004. Focal cortical dysplasias: surgical outcome in 67 patients in relation to histological subtypes and dual pathology. Brain 127:2406–2418.

Gibson EM, Purger D, Mount CW, Goldstein AK, Lin GL, Wood LS, Inema I, Miller SE, Bieri G, Zuchero JB, Barres BA, Woo PJ, Vogel H, Monje M. 2014. Neuronal Activity Promotes Oligodendrogenesis and Adaptive Myelination in the Mammalian Brain. Science 344:1252304.

Gruber V, Lang J, Endmayr V, Diehm R, Pimpel B, Glatter S, Anink JJ, Bongaarts A, Luinenburg MJ, Reinten RJ, Van Der Wel N, Larsen P, Hainfellner JA, Rössler K, Aronica E, Scholl T, Mühlebner A, Feucht M. 2021. Impaired myelin production due to an intrinsic failure of oligodendrocytes in mTORpathies. Neuropathol Appl Neurobio 47:812– 825.

Honke J, Hoffmann L, Coras R, Kobow K, Leu C, Pieper T, Hartlieb T, Bien CG, Woermann F, Cloppenborg T, Kalbhenn T, Gaballa A, Hamer H, Brandner S, Rössler K, Dörfler A, Rampp S, Lemke JR, Baldassari S, Baulac S, Lal D, Nürnberg P, Blümcke I. 2023. Deep histopathology genotype–phenotype analysis of focal cortical dysplasia type II differentiates between the GATOR1-altered autophagocytic subtype IIa and MTOR-altered migration deficient subtype IIb. Acta Neuropathol Commun 11:179.

Hornig J, Fröb F, Vogl MR, Hermans-Borgmeyer I, Tamm ER, Wegner M. 2013. The Transcription Factors Sox10 and Myrf Define an Essential Regulatory Network Module in Differentiating Oligodendrocytes. PLoS Genet 9:e1003907.

Kimura Y, Shioya A, Saito Y, Oitani Y, Shigemoto Y, Morimoto E, Suzuki F, Ikegaya N, Kimura Y, Iijima K, Takayama Y, Iwasaki M, Sasaki M, Sato N. 2019. Radiologic and Pathologic Features of the Transmantle Sign in Focal Cortical Dysplasia: The T1 Signal Is Useful for Differentiating Subtypes. AJNR Am J Neuroradiol 40:1060–1066.

Knowles JK, Xu H, Soane C, Batra A, Saucedo T, Frost E, Tam LT, Fraga D, Ni L, Villar K, Talmi S, Huguenard JR, Monje M. 2022. Maladaptive myelination promotes generalized epilepsy progression. Nat Neurosci 25:596–606.

Liewald D, Miller R, Logothetis N, Wagner H-J, Schüz A. 2014. Distribution of axon diameters in cortical white matter: an electron-microscopic study on three human brains and a macaque. Biol Cybern 108:541–557.

Lim JS, Kim W, Kang H-C, Kim SH, Park AH, Park EK, Cho Y-W, Kim S, Kim HM, Kim JA, Kim J, Rhee H, Kang S-G, Kim HD, Kim D, Kim D-S, Lee JH. 2015. Brain somatic mutations in MTOR cause focal cortical dysplasia type II leading to intractable epilepsy. Nat Med 21:395–400.

Lu QR, Sun T, Zhu Z, Ma N, Garcia M, Stiles CD, Rowitch DH. 2002. Common Developmental Requirement for Olig Function Indicates a Motor Neuron/Oligodendrocyte Connection. Cell 109:75–86.

Mei F, Wang H, Liu S, Niu J, Wang L, He Y, Etxeberria A, Chan JR, Xiao L. 2013. Stage-Specific Deletion of Olig2 Conveys Opposing Functions on Differentiation and Maturation of Oligodendrocytes. J Neurosci 33:8454–8462.

Meijer DH, Sun Y, Liu T, Kane MF, Alberta JA, Adelmant G, Kupp R, Marto JA, Rowitch DH, Nakatani Y, Stiles CD, Mehta S. 2014. An Amino Terminal Phosphorylation Motif Regulates Intranuclear Compartmentalization of Olig2 in Neural Progenitor Cells. J Neurosci 34:8507–8518.

Moreno-Jiménez EP, Flor-García M, Terreros-Roncal J, Rábano A, Cafini F, Pallas-Bazarra N, Ávila J, Llorens-Martín M. 2019. Adult hippocampal neurogenesis is abundant in neurologically healthy subjects and drops sharply in patients with Alzheimer’s disease. Nat Med 25:554–560.

Mühlebner A, Coras R, Kobow K, Feucht M, Czech T, Stefan H, Weigel D, Buchfelder M, Holthausen H, Pieper T, Kudernatsch M, Blümcke I. 2012. Neuropathologic measurements in focal cortical dysplasias: validation of the ILAE 2011 classification system and diagnostic implications for MRI. Acta Neuropathol 123:259–272.

Najm I, Lal D, Alonso Vanegas M, Cendes F, Lopes-Cendes I, Palmini A, Paglioli E, Sarnat HB, Walsh CA, Wiebe S, Aronica E, Baulac S, Coras R, Kobow K, Cross JH, Garbelli R, Holthausen H, Rössler K, Thom M, El-Osta A, Lee JH, Miyata H, Guerrini R, Piao Y, Zhou D, Blümcke I. 2022. The ILAE consensus classification of focal cortical dysplasia: An update proposed by an ad hoc task force of the ILAE diagnostic methods commission. Epilepsia 63:1899–1919.

Nave K-A, Werner HB. 2014. Myelination of the Nervous System: Mechanisms and Functions. Annu Rev Cell Dev Biol 30:503–533.

Nguyen LH, Mahadeo T, Bordey A. 2019. mTOR Hyperactivity Levels Influence the Severity of Epilepsy and Associated Neuropathology in an Experimental Model of Tuberous Sclerosis Complex and Focal Cortical Dysplasia. J Neurosci 39:2762–2773.

Palmini A, Andermann F, Olivier A, Tampieri D, Robitaille Y, Andermann E, Wright G. 1991. Focal neuronal migration disorders and intractable partial epilepsy: A study of 30 patients. Ann Neurol 30:741–749.

Rayatpour A, Farhangi S, Verdaguer E, Olloquequi J, Ureña J, Auladell C, Javan M. 2021. The Cross Talk between Underlying Mechanisms of Multiple Sclerosis and Epilepsy May Provide New Insights for More Efficient Therapies. Pharmaceuticals 14:1031.

Rushton WAH. 1951. A theory of the effects of fibre size in medullated nerve. J Physiol 115:101–122.

Scholl T, Mühlebner A, Ricken G, Gruber V, Fabing A, Samueli S, Gröppel G, Dorfer C, Czech T, Hainfellner JA, Prabowo AS, Reinten RJ, Hoogendijk L, Anink JJ, Aronica E, Feucht M. 2017. Impaired oligodendroglial turnover is associated with myelin pathology in focal cortical dysplasia and tuberous sclerosis complex. Brain Pathol 27:770–780.

Shepherd C, Liu J, Goc J, Martinian L, Jacques TS, Sisodiya SM, Thom M. 2013. A quantitative study of white matter hypomyelination and oligodendroglial maturation in focal cortical dysplasia type II. Epilepsia 54:898–908.

Sock E, Wegner M. 2021. Using the lineage determinants Olig2 and Sox10 to explore transcriptional regulation of oligodendrocyte development. Dev Neurobiol 81:892–901.

Srivastava A, Kumar K, Banerjee J, Tripathi M, Dubey V, Sharma D, Yadav N, Sharma MC, Lalwani S, Doddamani R, Chandra PS, Dixit AB. 2021. Transcriptomic profiling of high- and low-spiking regions reveals novel epileptogenic mechanisms in focal cortical dysplasia type II patients. Mol Brain 14:120.

Stassart RM, Möbius W, Nave K-A, Edgar JM. 2018. The Axon-Myelin Unit in Development and Degenerative Disease. Front Neurosci 12:467.

Steadman PE, Xia F, Ahmed M, Mocle AJ, Penning ARA, Geraghty AC, Steenland HW, Monje M, Josselyn SA, Frankland PW. 2020. Disruption of Oligodendrogenesis Impairs Memory Consolidation in Adult Mice. Neuron 105:150–164.e6.

Sternberger LA, Sternberger NH. 1983. Monoclonal antibodies distinguish phosphorylated and nonphosphorylated forms of neurofilaments in situ. Proc Natl Acad Sci USA 80:6126–6130.

Tomasi S, Caminiti R, Innocenti GM. 2012. Areal Differences in Diameter and Length of Corticofugal Projections. Cereb Cortex 22:1463–1472.

Tomassy GS, Berger DR, Chen H-H, Kasthuri N, Hayworth KJ, Vercelli A, Seung HS, Lichtman JW, Arlotta P. 2014. Distinct Profiles of Myelin Distribution Along Single Axons of Pyramidal Neurons in the Neocortex. Science 344:319–324.

Urbach H, Mast H, Egger K, Mader I. 2015. Presurgical MR Imaging in Epilepsy. Clin Neuroradiol 25:151–155.

Urbach H, Scheffler B, Heinrichsmeier T, Von Oertzen J, Kral T, Wellmer J, Schramm J, Wiestler OD, Blümcke I. 2002. Focal Cortical Dysplasia of Taylor’s Balloon Cell Type: A Clinicopathological Entity with Characteristic Neuroimaging and Histopathological Features, and Favorable Postsurgical Outcome. Epilepsia 43:33–40.

Van Tilborg E, Van Kammen CM, De Theije CGM, Van Meer MPA, Dijkhuizen RM, Nijboer CH. 2017. A quantitative method for microstructural analysis of myelinated axons in the injured rodent brain. Sci Rep 7:16492.

Vogel US, Thompson RJ. 1988. Molecular Structure, Localization, and Possible Functions of the Myelin-Associated Enzyme 2′,3′-Cyclic Nucleotide 3′-Phosphodiesterase. J Neurochem 50:1667–1677.

Wellmer J, Quesada CM, Rothe L, Elger CE, Bien CG, Urbach H. 2013. Proposal for a magnetic resonance imaging protocol for the detection of epileptogenic lesions at early outpatient stages. Epilepsia 54:1977– 1987.

Willard A, Antonic-Baker A, Chen Z, O’Brien TJ, Kwan P, Perucca P. 2022. Seizure Outcome After Surgery for MRI-Diagnosed Focal Cortical Dysplasia: A Systematic Review and Meta-analysis. Neurology [Internet] 98. Available from: https://www.neurology.org/doi/10.1212/WNL.0000000000013066

Zhao B, Zhang C, Wang X, Wang Y, Liu C, Mo J, Zheng Z, Zhang K, Shao X, Hu W, Zhang J. 2020. Sulcus-centered resection for focal cortical dysplasia type II: surgical techniques and outcomes. J Neurosurg135:266–272.

Zhou Q, Anderson DJ. 2002. The bHLH Transcription Factors OLIG2 and OLIG1 Couple Neuronal and Glial Subtype Specification. Cell 109:61– 73.

Zucca I, Milesi G, Medici V, Tassi L, Didato G, Cardinale F, Tringali G, Colombo N, Bramerio M, D’Incerti L, Freri E, Morbin M, Fugnanesi V, Figini M, Spreafico R, Garbelli R. 2016. Type II focal cortical dysplasia: Ex vivo 7 T magnetic resonance imaging abnormalities and histopathological comparisons. Ann Neurol 79:42–58.

