## Supplementary Table 1 for "Focal cortical dysplasia type II-dependent maladaptive myelination in the human frontal lobe"

Table 1. Clinical data of FCD IIa (A), FCD IIb (B) and controls (C) selected for this study.

A. FCD IIa cases

| **ID** | **Color code** | **Sex** | **Age at**  **epilepsy**  **onset (years)** | **Age at**  **surgery**  **(years)** | **Epilepsy**  **duration**  **(years)** | **Seizure**  **type** | **Seizure**  **frequency/**  **day** | **Neuro-**  **pathology** | **MRI findings** | **Brodmann**  **area** | **Methods**  **applied** |
| --- | --- | --- | --- | --- | --- | --- | --- | --- | --- | --- | --- |
| 1 |  | m | 0.2 | 6 | 5.8 | FA | n.a., during night | IIa | FCD | 8 | IHC, ISHH, PCR, ChIP |
| 2 |  | m | 8 | 22 | 14 | FA, FIA, BTCS | 2-3 | IIa | TM-FCD | 10 | IHC, ISHH, PCR, ChIP |
| 3 |  | f | 2 | 8 | 6 | FA | >1 | IIa | FCD | 8 | IHC, EM,  ISHH, PCR, ChIP |
| 4 |  | f | 18 | 23 | 5 | FA, FIA, BTCS | 0.07 | IIa | Neg. | 9 | IHC, EM,  ISHH, PCR, ChIP |
| 5 |  | f | 10 | 18 | 8 | FA, FIA | 8-10 | IIa | Neg. | 9 | IHC, EM,  ISHH, PCR, ChIP |
| 5 |  | f | 15 | 18 | 3 | FA, FIA, BTCS | 0.04 | IIa | FCD | n. a. | IHC, ISHH, PCR, ChIP |
| 7 |  | m | 0.6 | 10 | 9.4 | FA, FIA | 4.5 | IIa | FCD | 9 | IHC, ISHH, EM, PCR, ChIP |
| 8 |  | f | 1 | 8 | 7 | FIA | 0.36 | IIa | FCD | 9 | IHC, ISHH, PCR, PCR, ChIP |
| 9 |  | m | 4 | 38 | 34 | FA, FIA | 3 | IIa | Neg. | 44 | IHC, ISHH, PCR, PCR, ChIP |

B. FCD IIb cases

| **ID** | **Color code** | **Sex** | **Age at**  **epilepsy**  **onset (years)** | **Age at**  **surgery**  **(years)** | **Epilepsy**  **duration**  **(years)** | **Seizure**  **type** | **Seizure**  **frequency/**  **day** | **Neuro-**  **pathology** | **MRI findings** | **Brodmann**  **area** | **Methods applied** |
| --- | --- | --- | --- | --- | --- | --- | --- | --- | --- | --- | --- |
| 1 |  | m | 7.0 | 48 | 41.0 | FA, BTCS | 0.1,  during night | IIb | TM-FCD | 6 | IHC, EM,  ISHH, PCR, ChIP |
| 2 |  | m | 5 | 36 | 31 | FA, FIA | 6 | IIb | TM-FCD, | 8 | IHC, EM, ISHH, PCR, ChIP |
| 3 |  | f | 9 | 45 | 36 | FA, BTCS | 0.29 | IIb | TM-FCD | 6 | IHC, ISHH, PCR, ChIP |
| 4 |  | f | 9 | 27 | 18 | FA | 1 | IIb | TM-FCD | n.a. | IHC, ISHH,  PCR, ChIP |
| 5 |  | f | 12 | 40 | 28 | FA, BTCS | 1 | IIb | TM-FCD | 8 | IHC, EM, ISHH,  PCR, ChIP |
| 6 |  | m | 14 | 23 | 9 | FA, FIA, BTCS | 1 | IIb | TM-FCD | 8 | IHC, EM, ISHH,  PCR, ChIP |
| 7 |  | m | 9 | 38 | 29 | FA, BTCS | 1.2 | IIb | TM-FCD | 6 | IHC, ISHH, PCR, ChIP |
| 8 |  | f | 3 | 12 | 9 | FA, BTCS | multiple | IIb | TM-FCD | 6 | IHC, ISHH, PCR, ChIP |
| 9 |  | f | 16 | 34 | 18 | FIA, BTCS | 0.004 | IIb | TM-FCD | 44, 45 | IHC, ISHH, PCR, ChIP |

C. Control cases

| **ID** | **Color code** | **Sex** | **Age at**  **epilepsy**  **onset (years)** | **Age at**  **surgery**  **(years)** | **Epilepsy**  **duration**  **(years)** | **Seizure**  **type** | **Seizure**  **frequency/**  **day** | **MRI findings** | **Brodmann**  **area** | **Methods applied** |
| --- | --- | --- | --- | --- | --- | --- | --- | --- | --- | --- |
| 1 |  | f | 19 | 26 | 7 | FIA, BTCS | 0.43 | GG | 10 | IHC, EM, ISHH, ChIP, PCR |
| 2 |  | m | 12 | 17 | 5 | FIA, BTCS | 2-3 | Cystic lesion | 10 | IHC, EM,  ISHH, ChIP, PCR |
| 3 |  | m | 2 | 10 | 8 | FIA, BTCS | 2 | Cystic lesion | 9 | IHC, EM, ISHH, ChIP, PCR |
| 4 |  | f | 2 | 14 | 12 | FA | 1 | Blurring of GM-WM boundary | 11 | IHC, EM, ISHH, ChIP, PCR |
| 5 |  | m | 2 | 6.1 | 4.1 | FIA | 2 | Angiocentric glioma | 9 | ChIP, PCR |
| 6 |  | m | 9 | 26.7 | 17.7 | FA, BTCS | 0.3 | DNET | n.a. | IHC, ISHH, ChIP, PCR |
| 7 |  | f | 7 | 30 | 23 | FIA | n.a. | Ependymon | 10 | IHC, ISHH, ChIP, PCR |

ChIP, chromatin immunoprecipitation; BTCS: focal to bilateral tonic-clonic seizures; DNET: dysembryoplastic neuroepithelial tumor; EM, electron microscopy; FA: focal aware seizures; FCD, focal cortical dysplasia; FIA: focal impaired awareness seizures; GG, ganglioglioma; GM, gray matter; IHC, immunohistochemistry; ISHH, *in situ* hybridization histochemistry; n.a., not available; neg., negative; PCR, polymerase chain reaction; TM, transmantle sign; WM, white matter.
